# Transcription of Murine Endogenous Retrovirus MERVL Is Required for Progression of Development in Early Preimplantation Embryos

**DOI:** 10.1101/2022.07.20.500739

**Authors:** Akihiko Sakashita, Tomohiro Kitano, Hirotsugu Ishizu, Youjia Guo, Harumi Masuda, Masaru Ariura, Kensaku Murano, Haruhiko Siomi

## Abstract

Zygotic genome activation (ZGA) is a critical post-fertilization step that promotes totipotency and allows different cell fates to emerge in the developing embryo. MERVL, murine endogenous retrovirus-L, is transiently upregulated at the 2-cell stage around the time of ZGA. Although MERVL expression is widely used as a marker of totipotency, the role of this retrotransposon in mouse embryogenesis remains elusive. Here, we develop a method for consistent knockdown of interspersed copies of MERVL and show that MERVL RNA, but not encoded retroviral proteins and their long terminal repeat (LTR) promoter, is essential for accurate regulation of the host transcriptome and chromatin state during preimplantation development. MERVL knockdown (KD) results in embryonic lethality due to defects in differentiation and genomic stability. Furthermore, transcriptome and epigenome analysis revealed that MERVL-KD embryos retained an accessible chromatin state at, and aberrant expression of, a subset of 2-cell specific genes at mid-preimplantation stages. Taken together, our results suggest a model in which an endogenous retrovirus plays a critical role in regulating host cell fate potential.

## Introduction

Fertilization and early preimplantation development are processes in which unipotent gametes unite and acquire totipotency (Figure 1a). After fertilization, embryos undergo zygotic genome activation (ZGA), a process that is widely conserved in vertebrates ^1-3^. ZGA involves a transcriptional burst of hundreds-to-thousand of two-cell (2C)-specific genes. At this point gene expression switches from maternal mRNAs and proteins that were deposited in the oocyte to *de novo* transcripts expressed from the zygotic genome ^2,4^. ZGA occurs in two distinct waves called minor and major ZGA, respectively ^5^. In mice, minor ZGA occurs from S phase in the zygote to G1 phase in the early 2-cell stage embryo, whereas the major wave occurs during the second round of DNA replication at the middle-to-late 2-cell stage ^6,7^. Both waves of ZGA are critical for the embryo to acquire developmental competence. Indeed, inhibition of minor and major ZGA compromised preimplantation development at the 2-cell stage and later stages, respectively ^6,8^. However, the molecular events and chromatin reorganization that drive ZGA and lead to acquisition of totipotency and developmental competence are still enigmatic.

**Figure 1.**
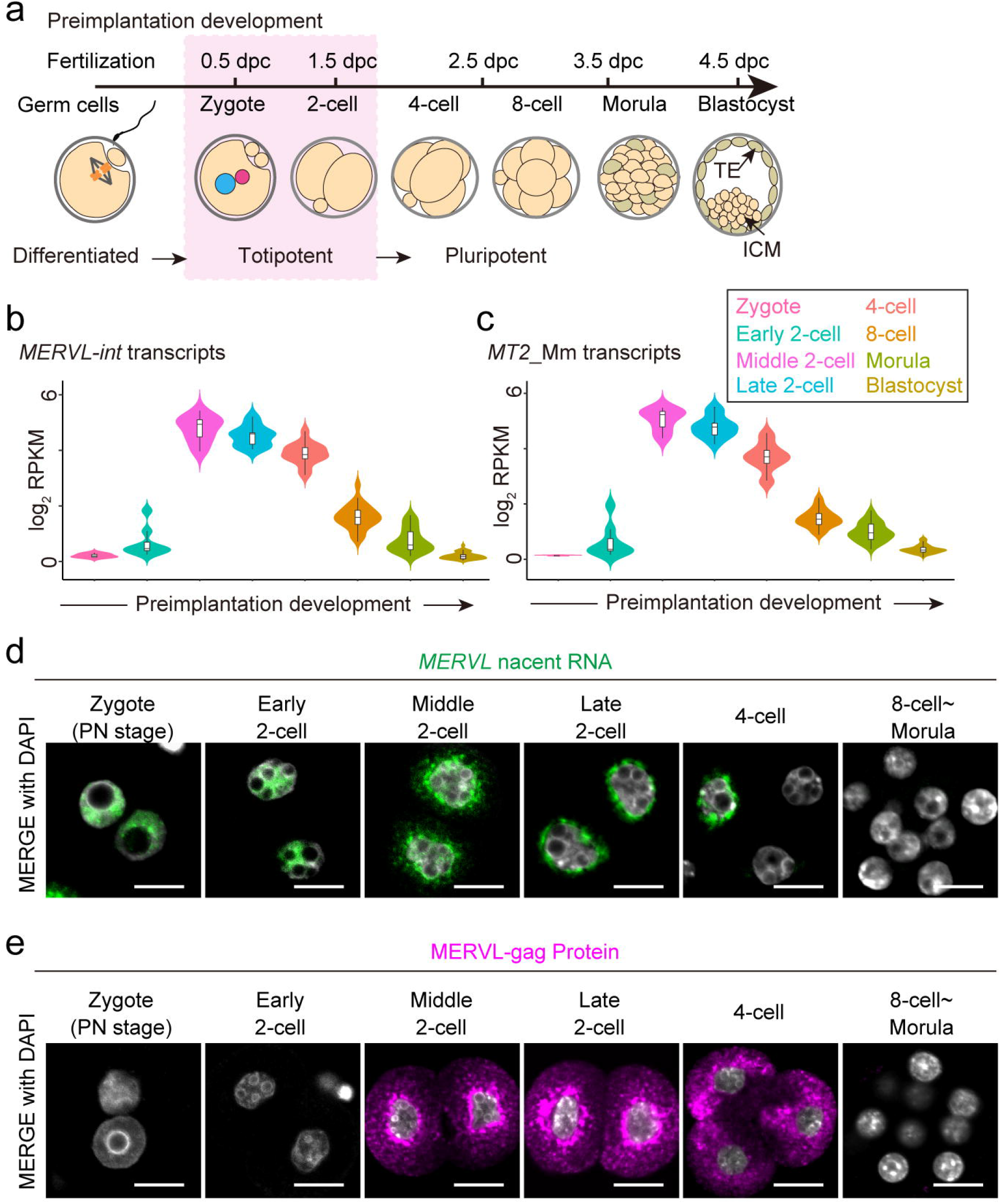
MERVL RNA exhibits dynamic nuclear-cytoplasmic expression during early stages of mouse preimplantation development. **(a)** Schematic of mouse preimplantation development. Totipotency is restricted to early-stage development (i.e. Zygote and 2-cell stages). The blastomere gradually transitions to a pluripotent state from the 4-cell stage onward and develops into a blastocyst consisting of inner cell mass (ICM) and trophectoderm (TE), prior to implantation in the uterus at 4.5-day post-coitum (dpc). **(b, c)** Violin plots showing the log_2_ expressions of *MERVL-int* **(b)** and its LTR promoter, *MT2_Mm* **(c)** during preimplantation development. **(d)** Representative images of smFISH with probes set for full-length MERVL RNA sequence with the DNA counterstained with DAPI during preimplantation development. Scale bar: 20 µm. **(e)** Representative images of immunofluorescence staining with antibody raised against the MERVL-gag protein with DNA counterstained with DAPI during preimplantation development. Scale bar: 20 µm.

Approximately 40% of the mouse genome is occupied by transposable elements (TEs), mobile genetic elements of which ∼10% are endogenous retrovirus (ERV) sequences ^9^. Notably, the expression of murine ERV with leucine transfer RNA primer (MERVL), is specifically activated at the 2-cell stage concomitant with ZGA ^10-12^. Recently, the transcription factor Dux, which is expressed during minor ZGA, was documented as an upstream regulator that activates 2C genes and MERVL ^13-15^. Furthermore, the MERVL long terminal repeat (LTR) promoter drives a subset of 2C genes and generates chimeric fusion transcripts linked to MERVL elements ^12^. The above findings suggest that Dux/MERV may activate an early transcriptional network that is required for ZGA and totipotency.

In 2012, Macfarlan and colleagues found that a rare transient cell population (∼1%) in mouse ESC and iPSC cultures expresses high levels of MERVL and 2C genes without expression of pluripotent inner cell mass (ICM) maker genes, such as *Oct4, Sox2* and *Nanog* ^12^. MERVL expression has been used as a marker for totipotent cells since MERVL+ cells from ESCs can commit to both embryonic and extraembryonic lineages after injection into recipient embryos at the 8-cell and morula stages ^12,16^. Subsequently, the regulatory axis by which MERVL activation and expanding cell potency has been reported; GATA-binding protein 2 (Gata2) directly binds to *MT2_Mm*, a LTR promoter of MERVL element, and promotes induction of MERVL and its proximal genes. In addition, depletion of *miR-34a*, a small non-coding RNA which directly inhibits *Gata2* expression at the post-transcriptional level in ESCs and iPSCs cultures, resulting in expansion of cell potency ^17^. Furthermore, the synergistic activation of *MT2_Mm* using the CRISPR activation system in ESCs drives the expression of adjacent 2C genes, which in turn expands cell potency ^16^. These observations in ESC and iPSC cultures suggest that MERVL may play a critical role in totipotency, differentiation and ontogeny in preimplantation embryos.

Despite of above findings, the function of MERVL itself remains unclear. This knowledge gap results from the technical hurdle that TEs, including MERVL, are present as hundreds or thousands of copies, preventing gene knockout using the TALEN or CRISPR/Cas9 systems ^18,19^. Only a few studies have attempted to determine the function of MERVL in preimplantation development using knockdown (KD) strategies. The results of these studies are inconsistent due to variability in targeting MERVL. Past studies that have examined MERVK-KD found either a subtle developmental delay or no obvious phenotype ^11,20,21^. In addition, the maternal and zygotic knockout of *Dux*, an upstream regulator of MERVL, was tolerated and ZGA and preimplantation development proceeded as normal ^22-25^. Given these reports, it has been assumed that MERVL expression is a consequence of global DNA demethylation and provides a marker of totipotency, but is not essential for preimplantation development ^26-28^.

Here, we overcome technical limitations in interrogating TE functions and analyze the role of MERVL in preimplantation development. To achieve this goal, we developed specific sets of antisense oligonucleotides (ASOs) targeting MERVL and successfully knocked down MERVL transcript. We confirmed that MERVL-KD embryos suffer a developmental delay and find that development halts at the morula-to-blastocyst transition. This phenotype is linked to major defects in differentiation and the failure of early lineage specification and genomic instability. We developed KD strategies that overcame previous barriers to analyzing MERVL function, and demonstrate that MERVL is essential for the progression of mouse preimplantation development. Our findings open up new avenues for investigation of TE function as well as critical events in early mammalian development.

## Results

### MERVL RNA exhibits distinct localization patterns during preimplantation stages of mouse development

Transcriptomic and immunocytochemical analysis of mouse preimplantation embryos have shown that MERVL is normally restricted to 2-cell stage embryos ^12,29^. To better understand the dynamics of MERVL expression, we first analyzed publicly available single-cell RNA-seq (scRNA-seq) datasets generated from each blastomere at 8 representative stages of preimplantation development ^30^ (Figure 1a). To define regions of non-redundant MERVLs in mouse genome, we used RepeatMasker annotation to annotate the genome for unique interspersed internal regions of MERVL (*MERVL-int*) and LTR promoters of the MERVL family (*MT2_Mm*). In this way, we filtered to 1,461 and 2,366 highly confident *MERVL-int* and *MT2_Mm* loci. Applying this annotation set to our RNA-seq processing pipeline, we detected the expression of individual MERVL copies in each blastomere from preimplantation embryos. Consistent with previous reports, the expression of MERVL and its LTR promoter culminated in the middle of the 2-cell stage and then gradually decreased until blastocyst stage (Figure 1b, c).

Next, we also set out to investigate the expression and localization of MERVL transcripts in preimplantation embryos using single-molecule fluorescence *in situ* hybridization (smFISH), an alternative approach to detect intrinsically low signal from each transcript and non-coding RNA, since the scRNA-seq dataset we used in this study only detects polyadenylated mRNA due to a methodological limitation. Interestingly, nascent RNA smFISH with a specific probe set for full-length MERVL revealed that MERVL expression is detectable in the zygote nucleus and early 2-cell stage embryos in which polyadenylated MERVL mRNA cannot be detected (Figure 1b, d). Nuclear MERVL RNA signals observed in the zygote and early 2-cell stage embryos gradually translocated into the cytoplasm from middle 2-cell stage onward. By the late 2-cell stage, MERVL was only detectable in the cytoplasm (Figure 1d). These changes in MERVL transcript localization were consistent with increased protein levels during the middle 2-cell stage, as determined by immunofluorescence staining with antibody raised against group-specific antigen (Gag) of MERVL, a major retroviral structural protein (Figure 1e). In summary, the expression of nuclear MERVL RNA was activated during minor ZGA. These observations raise the possibility that nuclear MERVL transcript has distinct roles in gene regulation in the early stages of preimplantation development compared to cytoplasmic MERVL RNA, leading us to investigate MERVL function further.

### MERVL knockdown results in defects in early lineage specification and embryonic lethality

Inconsistencies regarding the early embryonic phenotypes of MERVL-KD in previous studies ^11,20,21^, warrant further investigations. Hence, as a starting point for defining the role of MERVL, we reexamined the KD effects of MERVL on preimplantation development. To this end, we developed specific ASOs that target interspersed MERVL copies. Three independent ASOs target the coding regions of Gag and polymerase/reverse transcriptase (Pol) in full-length copies of MERVL (Figure 2a, Supplementary Figure S1a). After predicting the genome-wide target sites of individual ASOs using BLASTN, we confirmed 68.4% (n = 1,000/1,461) of highly confident MERVL copies were targeted by at least one ASO with a perfect match and 88.5% (n = 1,293/1,461) with ≤1 bp mismatch (Supplementary Figure S1b, c). Subsequently, we confirmed that our ASO sequences targeted full length MERVL (≥ 5 kb, n = 610), by combining three independent anti-MERVL ASOs (Supplementary Figure S1d). In addition, we experimentally validated the KD efficiency of each ASO using a recently developed ESC-based *in vitro* system (Supplementary Figure S1e and see Methods). After induction of ASOs into ESCs, MERVL expression was drastically reduced at both the mRNA and protein levels (Supplementary Figure S1f-h), corroborating that our ASOs efficiently targeted MERVL. In line with *in vitro* and computational analyses, injection of each ASO into the male pronucleus of zygotes leads to significant reduction of MERVL RNA signal at the late 2-cell stage (Supplementary Figure S2a). Since we noted that cocktail of three independent ASOs increased MERVL-KD efficiency (Figure 2b, Supplementary Figure S2b, c), mixed ASOs (1:1:1 = 20 µM) were used in subsequent experiments.

**Figure 2.**
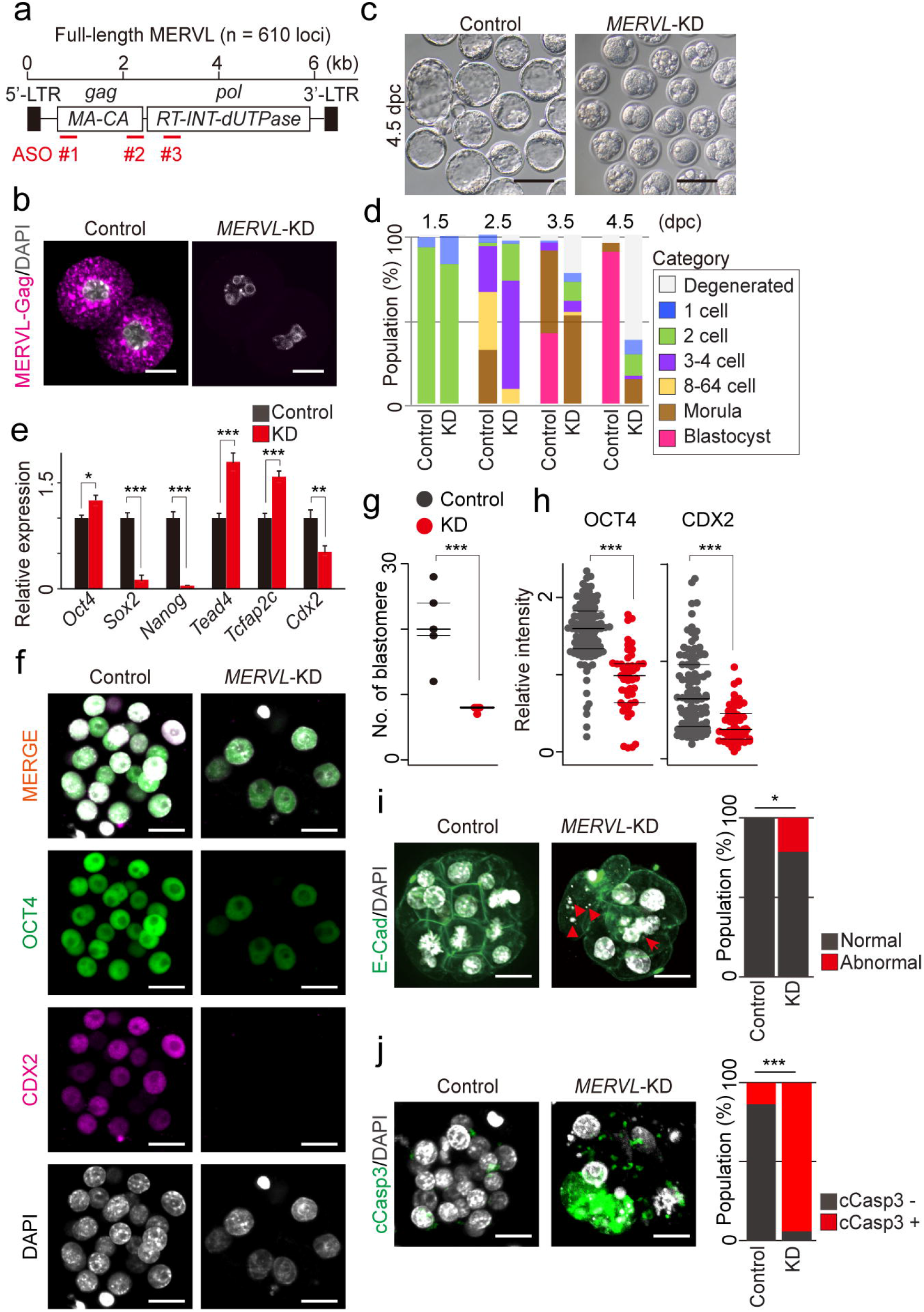
MERVL plays a critical role in early lineage specification and maintaining genomic stability of preimplantation embryos. **(a)** Schematic of MERVL indicating the positions of ASOs. Intact MERVL encodes the retroviral proteins; Gag (Group-specific antigens, comprised of MA, matrix; CA, capsid proteins) and Pol (Polymerase, comprised of RT, reverse transcriptase; INT, integrase; dUTPase, dUTP phosphatase). **(b)** Representative images of immunofluorescence staining with antibody raised against MERVL-gag protein with DNA counterstained with DAPI in control and MERVL-KD 2-cell embryos at 1.5 dpc. Scale bar: 20 µm. **(c)** Representative phase-contrast Images of 4.5 dpc blastocysts in control and MERVL-KD. Scale bar: 100 µm. **(d)** Percentage of embryos by stages of development at 1.5, 2.5, 3.5 and 4.5 dpc in control and MERVL-KD. **(e)** Expressions of ICM- and trophectoderm-associated genes (Oct4, Sox2 and Nanog are expressed in ICM; Tead4, Tcfap2c and Cdx2 are expressed in trophectoderm) as measured by quantitative RT-PCR in control and MERVL-KD morula at 3.5 dpc. Relative gene expression is quantified with ΔΔCt method and normalized to *Actb* expression. Error bars represent mean± s.e.m.: *P<0.05, **P<0.01, ***P<0.001, two-tailed unpaired t-tests. **(f)** Representative images of immunofluorescence staining with antibodies raised against OCT4 and CDX2 with the DNA counterstained with DAPI in control and MERVL-KD morula at 3.5 dpc. Scale bar: 20 µm. **(g, h)** Dot plots showing the total number of blastomere per embryo and relative intensity for OCT4 and CDX2 per nucleus in control and MERVL-KD morula at 3.5 dpc, normalized to DAPI signal. ***P<0.001, two-tailed unpaired t-tests. **(i)** Left: Representative images of immunofluorescence staining with E-cadherin (E-Cad) antibody. DNA is counterstained with DAPI in control and MERVL-KD morula at 3.5 dpc. Arrows and arrowheads indicate nuclear deformation and micronuclei, signs of genome instability, respectively. Scale bar: 20 µm. Right: Bar chart showing the percentage of abnormal nuclear morphology at morula stage, as determined by DAPI staining. *P<0.05, chi-square test. **(j)** Left: Representative images of immunofluorescence staining with antibody raised against cleaved Caspase 3 (cCasp3) with the DNA counterstained with DAPI in control and MERVL-KD morula at 3.5 dpc. Right: Bar chart showing the percentage of apoptotic cells per embryo at morula stage. ***P<0.001, chi-square test.

Next, we monitored the effects of MERVL-KD using ASOs on preimplantation development (Figure 2c, d). The rate of development of 2-cell embryos upon MERVL-KD was comparable to that of controls : 90.3% for the control ASO and 83.4% for MERVL-KD (*P*=0.172, chi-square test, Figure 2d). However, MERVL-KD embryos displayed significant a developmental delay from 2.5 day post-coitum (dpc). In agreement, despite longer culture, a greater percentage of MERVL-KD embryos remained at the 2-to 4-cell stage: 30.9% for control and 91.1% for MERVL-KD at 2.5 dpc (\*\**P*=0.00533, chi-square test, Figure 2d). Strikingly, at 4.5 dpc, when viable normal embryos should have reached the blastocyst stage, almost no blastocyst formation was observed in MERVL-KD embryos (Figure 2c,d). Overall, these findings revealed that majority of MERVL-KD embryos suffered a developmental delay and halted development at morula-to-blastocyst transition.

Of note, the phenotype of MERVL-KD embryos is reminiscent of that observed in loss-of-function of genes associated with ICM and trophectoderm differentiation ^31-35^, implying that MERVL transcription might function in early cell lineage specification during preimplantation development. To test this hypothesis, we examined mRNA levels of genes related to ICM (*Oct4, Sox2* and *Nanog*) and trophectoderm differentiation (*Tead4, Tcfap2c* and *Cdx2*) in MERVL-KD morula-like embryos at 3.5 dpc, using quantitative RT-PCR (Figure 2e). *Sox2, Nanog* and *Cdx2* mRNA levels were significantly reduced in MERVL-KD morula, in contrast to *Oct4, Tead4* and *Tcfap2c* transcripts which increased (Figure 2e). This data suggests that MERVL transcription during early stages of preimplantation development is required for subsequent proper expression of genes linked to the earliest cell lineage specification events. To further investigate the molecular etiologies of the phenotype upon MERVL-KD, we examined OCT4 and CDX2 protein expression using immunofluorescence analysis at the morula stage (Figure 2f). MERVL-KD morula is notably comprised of significantly fewer blastomeres, although MERVL-KD blastomeres apparently underwent compaction similar to those in control embryos (Figure 2g). Moreover, we observed a significant reduction in OCT4 and CDX2 protein expression (Figure 2h). Notably, MERVL-KD embryos also displayed increased hallmarks of genomic instability such as nuclear deformation and micronuclei, as visualized by DAPI staining (Figure 2i), suggesting that MERVL is linked to cell death and reduction in numbers of blastomere. Indeed, we confirmed that the population of apoptotic cells significantly increased in MERVL-KD embryos (Figure 2j). In sum, the depletion of MERVL transcript results in disruption of lineage specification, cell death, and ultimately early embryonic lethality.

### *Cis*-acting nascent RNA and/or MERVL transcription is essential for normal preimplantation development

Since some of interspersed MERVL copies with intact ORF encode functional proteins for retrotransposition (Figure 2a, Supplementary Figure S1a, b), transcribed MERVL RNA is stochastically processed and translated to retroviral proteins by host machinery ^36,37^. Therefore, we next interrogated which products from MERVL : *cis*-acting, *trans*-acting RNA or retroviral proteins, carried the functional essence required for preimplantation development. In order to assess the functionality of retroviral proteins, we utilized two independent siRNAs to KD MERVL, instead of ASOs. ASO-mediated silencing occurs via nuclear RNase H while siRNAs are loaded onto cytoplasmically localized Argonaute proteins to form RNA-induced silencing complexes (RISCs) that cleave their RNA targets, thus preferentially suppressing protein translation (Figure 3a)^38^. The day following siRNA injection, no MERVL-Gag protein was detected in late 2-cell stage embryos upon siRNA-mediated KD, confirming that siRNA-mediated silencing efficiently suppressed translation of MERVL retroviral protein (Figure 3b). Consequently, although the phenotypes of siRNA-mediated MERVL-KD are inconsistent between previous studies ^20,21^, we confirmed that siRNA-mediated KD of MERVL had no significant impact on preimplantation development. No changes in developmental rate were observed and MERVL-KD embryos reached the blastocyst stage, similar to control embryos : 65.7% for control and 70.0% for MERVL-KD (*P*=0.712, chi-square test). There was no evidence of developmental delay and morphological defects (Figure 3c, d). Retroviral proteins encoded by MERVL are therefore dispensable for preimplantation development.

**Figure 3.**
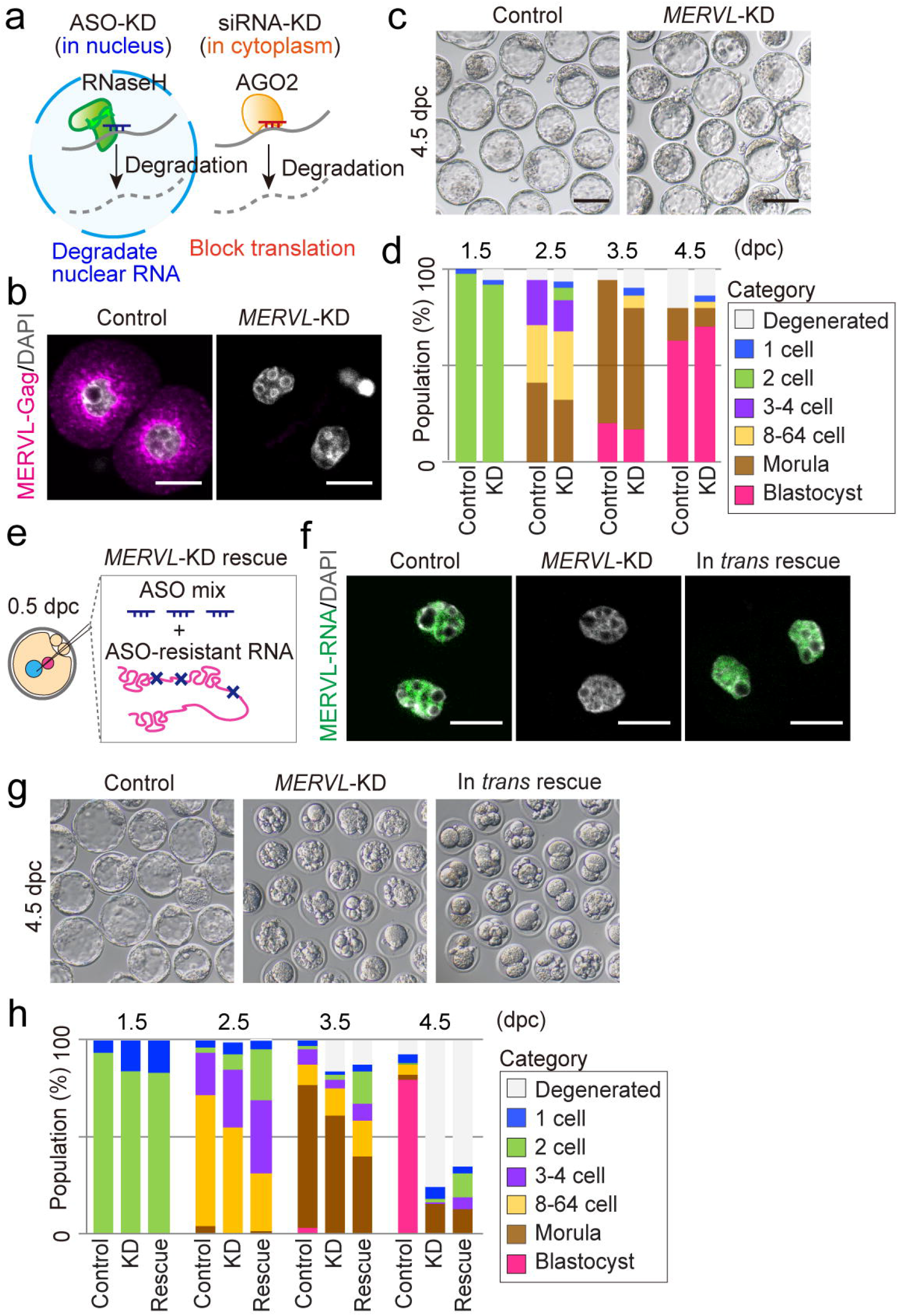
Retroviral proteins and trans-acting MERVL RNA are dispensable for early preimplantation development. **(a)** Schematic of the distinct mechanisms underlying ASO vs. siRNA-mediated RNA targeting. **(b)** Representative images of immunofluorescence staining with antibody raised against MERVL-gag protein with the DNA counterstained with DAPI in control and siRNA-mediated MERVL-KD 2-cell embryos at 1.5 dpc. Scale bar: 20 µm. **(c)** Representative phase-contrast Images of 4.5 dpc blastocysts in control and MERVL-KD by siRNA. Scale bar: 100 µm. **(d)** Percentage of embryos by developmental stage at 1.5, 2.5, 3.5 and 4.5 dpc in control and MERVL-KD by siRNA. **(e)** Schematic of experimental procedure for *in trans* rescuing MERVL-KD by ASOs. Transcribed ASO-resistant MERVL-RNA is co-injected into the male pronucleus of 0.5 dpc zygotes with MERVL ASOs. **(f)** Representative images of smFISH with probe specific to the full-length MERVL RNA sequence. DNA is counterstained with DAPI in early 2-cell embryos in each experimental condition. Scale bar: 20 µm. **(g)** Representative phase-contrast images of 4.5 dpc blastocysts in each experimental condition. Scale bar: 100 µm. **(h)** Percentage of embryos by developmental stage at 1.5, 2.5, 3.5 and 4.5 dpc in each experimental condition.

Given the above data, we next focused our analysis on functions of the MERVL RNA acting in *cis-* or in *trans-*. To determine whether transcribed MERVL RNA plays a critical role in preimplantation development, we utilized a MERVL RNA construct with point mutations conferring resistance to ASO-mediated KD (Figure 3e). In this rescue experiment we co-injected an ASO-resistant MERVL RNA with mixed ASOs into zygotes. Nascent MERVL RNA signal was observed in nuclei from MERVL-KD early 2-cell stage embryos at levels comparable to control embryos when ASO-resistant MERVL RNA was present, as determined by smFISH with specific probe set for full-length MERVL (Figure 3f). However, the embryonic lethal phenotype was not rescued by co-induction of trans-acting ASO-resistant MERVL RNA with MERVL-KD ASOs (Figure 3g, h). At day 4 after the co-injection, no blastocysts were also observed upon induction of ASO-resistant MERVL RNA along with ASO-mediated KD, as well as in MERVL-KD embryos (Figure 3g, h). Our results revealed that the exogenously expressed (trans-acting) MERVL transcript could not rescue endogenous MERVL knockdown, suggesting that the complete rescue may require either cis-acting RNA or transcription itself, which are driven by MERVL loci. Collectively, our data demonstrated that neither encoded retroviral proteins nor *trans*-acting RNA of MERVL is indispensable for preimplantation development.

### The expression of a subset of 2C genes is dysregulated in ASO-mediated MERVL-KD preimplantation embryos

The data indicate that expression of interspersed MERVL copies is coordinated with the minor and major wave of ZGA at the zygote and 2-cell stage (Figure 1)^6^. Therefore, we next assessed the effects of ASO-mediated MERVL-KD on ZGA. We first made use of 5’-ethynyluridine (EU) to assay global transcriptional levels in control and MERVL-KD 2-cell stage embryos (Figure 4a). We observed no significant change in zygotic transcriptional activity between control and MERVL-KD 2-cell stage embryos during major ZGA (Figure 4b, c).

**Figure 4.**
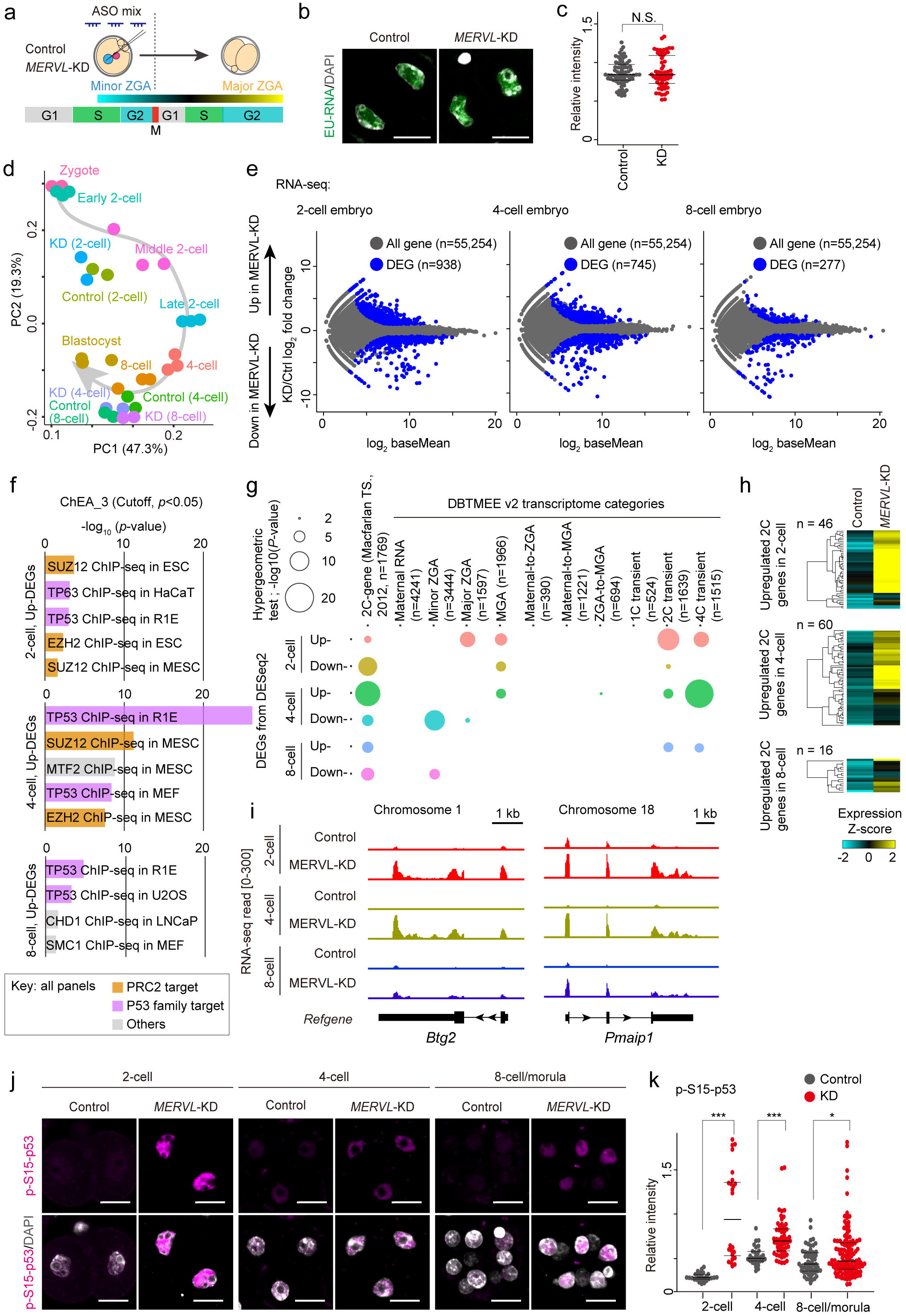
Ablation of MERVL impacts the expression of a subset of 2C-specific gene through preimplantation development. **(a)** Schematic of timing for minor and major ZGA (adapted from ^6^). **(b)** Representative images of EU incorporation assay with the DNA counterstained with DAPI in control and MERVL-KD 2-cell stage embryos. Scale bar: 20 µm. **(c)** The relative fluorescence intensity of EU incorporation per nucleus, normalized to DAPI signal. N.S., not significant; two-tailed unpaired t-tests. **(d)** Bidimensional principal component analysis of gene expression profiles in control and MERVL-KD embryos through preimplantation development. **(e)** RNA-seq differential gene expression analysis: MERVL-KD versus control embryos obtained at 2-cell, 4-cell and 8-cell stages. 938, 745 and 277 genes evinced significant changes in expression in MERVL-KD embryos (blue circle, FoldChange≥|2|, *Padj*<0.05; binomial test with Benjamini–Hochberg correction). **(f)** Predicted factors that are upstream of all up-regulated genes upon MERVL-KD, assessed by the ChEA database. The enrichments of known target genes of the polycomb repressive complex 2 (PRC2) and p53 family are highlighted in orange and purple. **(g)** Bubble plot shows overlap between all DEGs with the list of 2C genes and DBTMEE v2 transcriptome categories. The bubble plot sizes show the –log_10_[P values] derived from a hypergeometric test. **(h)** Heatmaps showing expression levels of 2C genes overlapped with upregulated DEGs in MERVL-KD embryos at 2-cell (n = 46), 4-cell (n =60) and 8-cell stages (n = 16). Expression level of each gene is shown in Z-score, calculated by subtracting the mean expression value and dividing by standard deviation. **(i)** Track views show RNA-seq signals in control and MERVL-KD embryos at 2-cell, 4-cell and 8-cell stages, on two representative 2C-gene loci (as defined in Macfarlan TS. et al. Nature, 2012). **(j)** Representative images of immunofluorescence staining with antibody raised against phosphorylated p53 on serine 15 (p-S15-p53) with the DNA counterstained with DAPI in control and MERVL-KD embryos at 2-cell, 4-cell and 8-cell stages. Scale bar: 20 µm. **(k)** Dot plots showing the relative intensity for p-S15-p53 per nucleus in control and MERVL-KD embryos at 2-cell, 4-cell and 8-cell stages, normalized to DAPI signal. *P<0.05, ***P<0.001, two-tailed unpaired t-tests.

To further investigate the molecular mechanism that underlies the developmental arrest and embryonic lethality in ASO-mediated MERVL-KD embryos, we carried out total RNA-seq analyses with depletion of ribosomal RNA, for 2-cell, 4-cell and 8-cell stage embryos upon control and MERVL-KD conditions (Supplementary Figure S3a, b). Total RNA-seq dataset provided accurate gene expression profiles; a summarized list is shown in Supplementary DataSet1 and 2. In line with principal component analysis (PCA) with published RNA-seq data in preimplantation embryos ^30^, our RNA-seq data accurately represent the transcriptome at each stage in preimplantation embryos. Transcriptional profiles of MERVL-KD embryos at each stage were not clearly distinguishable from those of control embryos (Figure 4d).

However, when we compared differentially expressed genes (DEGs) between control and MERVL-KD embryos with DESeq2, we found that 938, 745 and 277 genes (of manually annotated in Gencode (vM25) : n = 55,254) were significantly dysregulated in MERVL-KD embryos, at the 2-cell, 4-cell and 8-cell stages, respectively (fold change≥|2|, adjacent *P* value (*Padj*) < 0.01, Figure 4e, Supplementary DataSet3). In contrast to annotated genes, among all interspersed repetitive elements, only MERVL and a small number of closely related subtypes were significantly downregulated in MERVL-KD pre-implantation embryos (Supplementary Figure S4a, b), suggesting that KD of MERVL had no significant impact on expression of other types of repetitive elements, such as non-LTR type retrotransposons and satellite repeats.

Since the LTR promoter of MERVL (*MT2_Mm*) has been reported to drive chimeric fusion transcripts with adjacent target genes that are specific to the 2-cell stage ^12^, we also investigated the occurrence and expression level of chimeric fusion transcripts, using our paired-end RNA-seq reads from control and MERVL-KD 2-cell stage embryos (Supplementary Figure S4c). Largely consistent with previous reports (Macfarlan *et al*., 2012), we detected 77 genes that generated chimeric fusion transcripts with junction to *MT2_Mm* on the exon regions in both control and MERVL-KD embryos. However, the overall expression level of chimeric fusion transcripts is unchanged in MERVL-KD embryos (Supplementary Figure S4c).

As described above, we identified DEGs during preimplantation development of MERVL-KD embryos. To better understand the biological function of the identified DEGs, we performed gene ontology (GO) and pathway analyses on each set of DEGs. DEGs that are upregulated in MERVL-KD embryos are enriched in genes associated with “apoptosis/cell death” and “cell cycle”. This contrasts with downregulated DEGs which are preferentially comprised of genes associated with “DNA repair in response to DNA damage” (Supplementary Figure S3c). Interestingly, the ChIP Enrichment Analysis (ChEA) web tool ^39^ indicate that polycomb repressive complex 2 (PRC2)- and TP53 (also known as p53)-target genes were significantly enriched in each set of upregulated DEGs (Figure 4f). No significant enrichment was observed in each set of downregulated DEGs. Taken together, these transcriptome analyses in MERVL-KD embryos raise the possibility that DNA damage and activated p53 ectopically accumulate in the nuclei of MERVL-KD embryos which in turn leads to embryonic lethality via apoptosis.

Since DNA damage and activated p53 contribute to the induction of 2C genes in early stage preimplantation embryos and 2-cell like cells (2CLC) ^40^, we reasoned that the genes significantly up-regulated in MERVL-KD embryos might be enriched for 2C and p53-target genes. To test this hypothesis, we next compared DEGs with a list of 2C genes (n = 1,769, as defined in ^12^) and a predefined transcriptomic atlas of early mouse embryogenesis (DBTMEE_v2)^41,42^. Significant enrichment of 2C genes was observed in all DEG sets during preimplantation development (*P*<0.01, hypergeometric test for overrepresentation, Figure 4g). Strikingly, MERVL-KD 4-cell stage embryos exhibited the highest increase in 2C gene transcript levels (Figure 4g, h). In addition, 2C- and 4C-transient genes were also enriched in all sets of upregulated DEGs (Figure 4g). These findings argue that MERVL-KD embryos still retain a 2C/totipotent-like transcriptome even at mid-preimplantation stages (i.e. 4-cell and 8-cell stages). Indeed, representative track views demonstrated that 2C gene loci (e.g. *Btg2* and *Pmaip1*) maintain abnormally high expression levels of 2C genes in MERVL-KD preimplantation embryos. In contrast, 2C gene expression is restricted to the 2-cell stage in control embryos (Figure 4i). To determine whether ectopic expression of 2C genes in MERVL embryos is correlated with p53 activity, we performed immunofluorescence analysis with antibody against phospho-S15-p53 in control and MERVL-KD embryos (Figure 4j). Consistent with our hypothesis, we observed a significant increase in phospho-S15-p53 signal intensity in all developmental stages of MERVL-KD embryos, compared to controls (Figure 4k). Together with the embryonic phenotype in siRNA-mediated KD (Figure 3), these results indicate that a subset of 2C genes are persistently active due to defects in MERVL transcription and/or *cis*-acting RNA.

### MERVL modulates dynamic changes in accessible chromatin during the transition phase from totipotency to pluripotency

The totipotent-to-pluripotent transition in preimplantation embryos coincides with changes in chromatin mobility. Indeed, FRAP of core histones at the 2-cell stage indicates that higher chromatin mobility characterizes the 2-cell stage, but that mobility significantly decreases as development proceeds ^43^. Hereby, to further investigate the correlation between transcription and chromatin state ^44^, we next asked whether MERVL-KD impacts chromatin accessibility at the transcription start sites of dysregulated 2C genes. To investigate differences in genome-wide chromatin organization between control and MERVL-KD embryos at nucleosomal resolution, we made use of the optimized miniATAC-seq method (Supplementary DataSet1) ^45^. We used this approach due to the limited numbers of available MERVL-KD embryos (Supplementary Figure S5a). Two independent biological replicates confirmed that ATAC-seq signals are consistently detected and are highly correlated between replicates (Supplementary Figure S5b,c). Biological replicates were pooled for downstream analyses after confirming reproducibility. Akin to the total RNA-seq analysis (Figure 4d), the overall enrichment pattern of ATAC-seq signals at 10 kb intervals across the genome was modestly changed in MERVL-KD embryos during preimplantation development (Figure 5a). Meanwhile, open chromatin at MERVL loci was significantly reduced in MERVL-KD 2-cell stages embryos (Figure 5b). Since ASO-mediated KD might trigger premature transcriptional termination via degradation of the residual Pol II-associated transcripts ^46^, we suggest that the reduction of chromatin openness at MERVL loci is associated with premature termination of MERVL transcripts induced by ASO-mediated KD.

**Figure 5.**
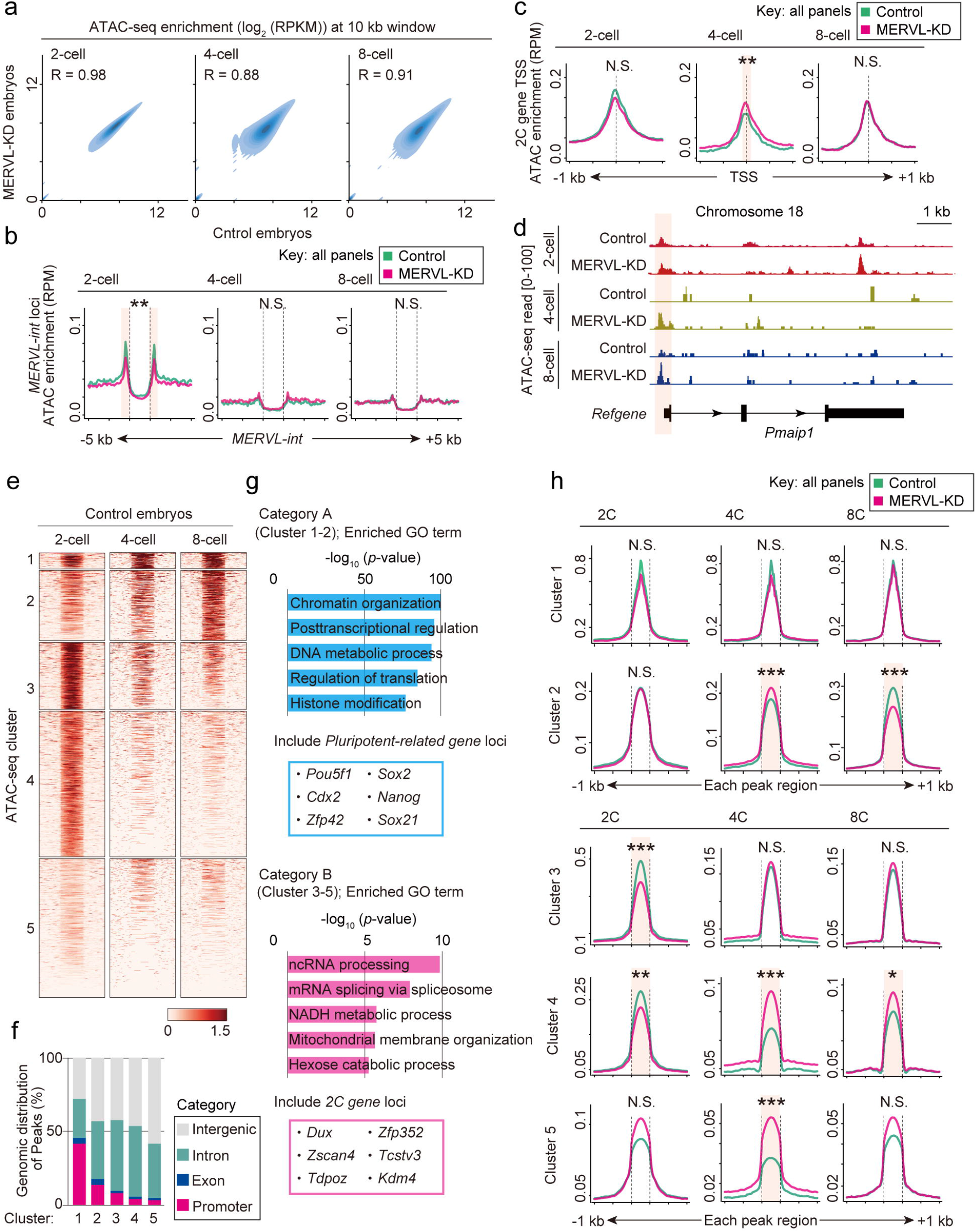
MERVL-KD embryos retain 2C-like chromatin accessibility even at mid-preimplantation stage. **(a)** Genome-wide correlation of ATAC-seq signals by stages of development between control and MERVL-KD embryos. Enrichment levels per 10 kb bin are shown in log_2_RPKM values. The Pearson correlation coefficient values (R) indicate the similarity between control and MERVL-KD embryos. **(b)** Average tag density plots of ATAC-seq enrichment around MERVL-int copies (±5 kb) by stages of development in control and MERVL-KD embryos. \*\**Padj*<0.01; N.S., not significant; Mann-Whitney U-test with Bonferroni correction. **(c)** Average tag density plots of ATAC-seq enrichment around TSS (±1 kb) of 2C genes by stages of development in control and MERVL-KD embryos. \*\**Padj*<0.01; N.S., not significant; Mann-Whitney U-test with Bonferroni correction. **(d)** Track view shows ATAC-seq signals in control and MERVL-KD embryos at 2-cell, 4-cell and 8-cell stages, on representative 2C-gene locus (as defined in Macfarlan *et al*. 2012). **(e)** Heatmaps showing ATAC-seq signals across all peak regions (±1 kb, identified with MACS2; *P*<1×10^−5^) in control embryos at 2-cell, 4-cell, and 8-cell stages. Each peak was ordered by k-means clustering of ATAC-seq signal and yielded five clusters. Cluster 1-2, increased accessibility during preimplantation development (defined as Category A); Cluster 3-5, were transiently accessible at 2-cell stage or reduced accessibility during preimplantation development (defined as Category B). **(f)** Stacked bar chart shows ATAC-seq peak distributions across genomic entities (intergenic, intron, exon and promoter) in each cluster. **(g)** Gene ontology analysis of genes adjacent to ATAC-seq peaks of Cluster1-2 (Category A) and Cluster3-5 (Category B). The pluripotent-related and 2C genes were obviously enriched in gene set adjacent to Category A and Category B, respectively. **(h)** Average tag density plots of ATAC-seq enrichment around peak regions (±1 kb) in each cluster by stages of development in control and MERVL-KD embryos. \**Padj*<0.05; \*\**Padj*<0.01; \*\*\**Padj*<0.001; N.S., not significant; Mann-Whitney U-test with Bonferroni correction.

Next, we determined whether the transcriptional changes in MERVL-KD embryos was associated with their chromatin states by examining enrichment of ATAC-seq signals at the transcription start sites (TSSs) of 2C genes whose expression was dysregulated in MERVL-KD preimplantation embryos. ATAC-seq signals showed significantly higher enrichment at the TSSs of 2C genes that retain high expression in MERVL-KD embryos at the 4-cell stage compared to control embryos (Figure 5c). In line with this finding, a representative track view showed that an accessible chromatin state was retained at the TSS of *Pmaip1*, a 2C gene. This open state persisted into the mid-preimplantation stage (i.e. 4-cell and 8-cell embryos) upon MERVL-KD, a time when this locus is repressed/closed in control embryos (Figure 5d). These results confirm that ectopic expression of 2C genes in mid-preimplantation embryos upon MERVL-KD is associated with accessible chromatin states at these loci.

The above findings raised the possibility that open chromatin structure of genes that are specific to totipotent state, was changed to inaccessible state in a MERVL-dependent manner as development proceeds. To address this possibility, we first characterized genome-wide chromatin accessibility states during preimplantation development, using ATAC-seq data from control embryos. k-means clustering on normalized ATAC-seq signals in peak regions yielded five clusters that fall into two distinct categories by peak region. Category A became accessible gradually as development proceeds (comprised of cluster 1 (n = 8,691) and 2 (n = 37,735), t) and Category B genes were specifically accessible in 2-cell stage embryos but exhibited loss of accessibility progressively or shortly after the 2-cell stage (comprised of cluster 3 (n =35,948), 4 (n = 78,297) and 5 (n= 74,556)) (Figure 5e). Among each cluster, approximately half of ATAC-seq peaks were enriched at largely genic regions, including promoter, exon and intronic regions (Figure 5f). GO analysis revealed that genes adjacent to peaks in Category A were highly enriched for roles in “chromatin organization” and “posttranscriptional regulation”, such as those involved in regulation of the pluripotent state (Figure 5g; top) ^47,48^. Moreover, the peaks in Category A contained genic regions that associated with pluripotency including *Pou5f1/Oct4* and *Cdx2* (Figure 5g; top, Supplementary DataSet4). Conversely, the peaks in Category B were characterized by genes and biological terms associated with totipotent- and 2C states, such as *Dux* and *Zscan4c* whose expression was restricted to 2-cell stage embryos (Figure 5g; bottom, Supplementary DataSet4)^13,14,49^. These observations revealed that dynamic changes in chromatin accessibility that occur shortly after 2-cell stage may be important for the transition from totipotency to pluripotency. Finally, to test the hypothesis that MERVL transcription contributes to changes in chromatin accessibility at the transition from the totipotent to pluripotent state, we examined ATAC-seq signals in each cluster and compared control and MERVL-KD embryos. In Category B (cluster 3-5), analysis of average tag densities of ATAC-seq signals confirmed that chromatin accessibility was significantly decreased in peaks of both cluster 3 and 4 in MERVL-KD 2-cell stage embryos (Figure 5h). In contrast, from 4-cell stage onward, ATAC-seq signals were significantly increased in peaks of cluster4 in MERVL-KD embryos (Figure 5h), suggesting that a 2C-like chromatin state was partially retained during preimplantation development of MERVL-KD embryos. In addition, among Category A (cluster 1-2) genes, which are related to cell fate transition and pluripotency, we observed a significant reduction of ATAC-seq signals in cluster 2 peaks in MERVL-KD 8-cell stage embryos (Figure 5h). This reduction may be associated with defects in early lineage specification upon MERVL-KD (Figure 2e-h). Taken together, these results support the proposal that MERVL has critical roles in chromatin remodeling and in differentiation from the totipotent to the pluripotent state.

## Discussion

In summary, we provided functional evidence that transcriptional activation of the mouse-specific endogenous retrovirus, MERVL, is essential for progression of development in mouse preimplantation embryos. We efficiently depleted MERVL RNA and encoded retroviral proteins by developing an optimized KD system and demonstrated that KD of MERVL RNA (but not encoded retroviral protein) results in embryonic lethality with profound defects in development and is associated with retaining a 2C-like transcriptome and chromatin state (Figure 6). These findings suggest the possibility that MERVL transcription in totipotent cells may act as the switch for the transition from totipotency to pluripotency and is responsible for the onset of differentiation and ontogeny (Figure 6). Although the precise molecular mechanism underlying dysregulation of 2C genes upon MERVL-KD still remains elusive, we provide new insight into the interaction between transposable elements and their hosts in the regulation of totipotency.

**Figure 6.**
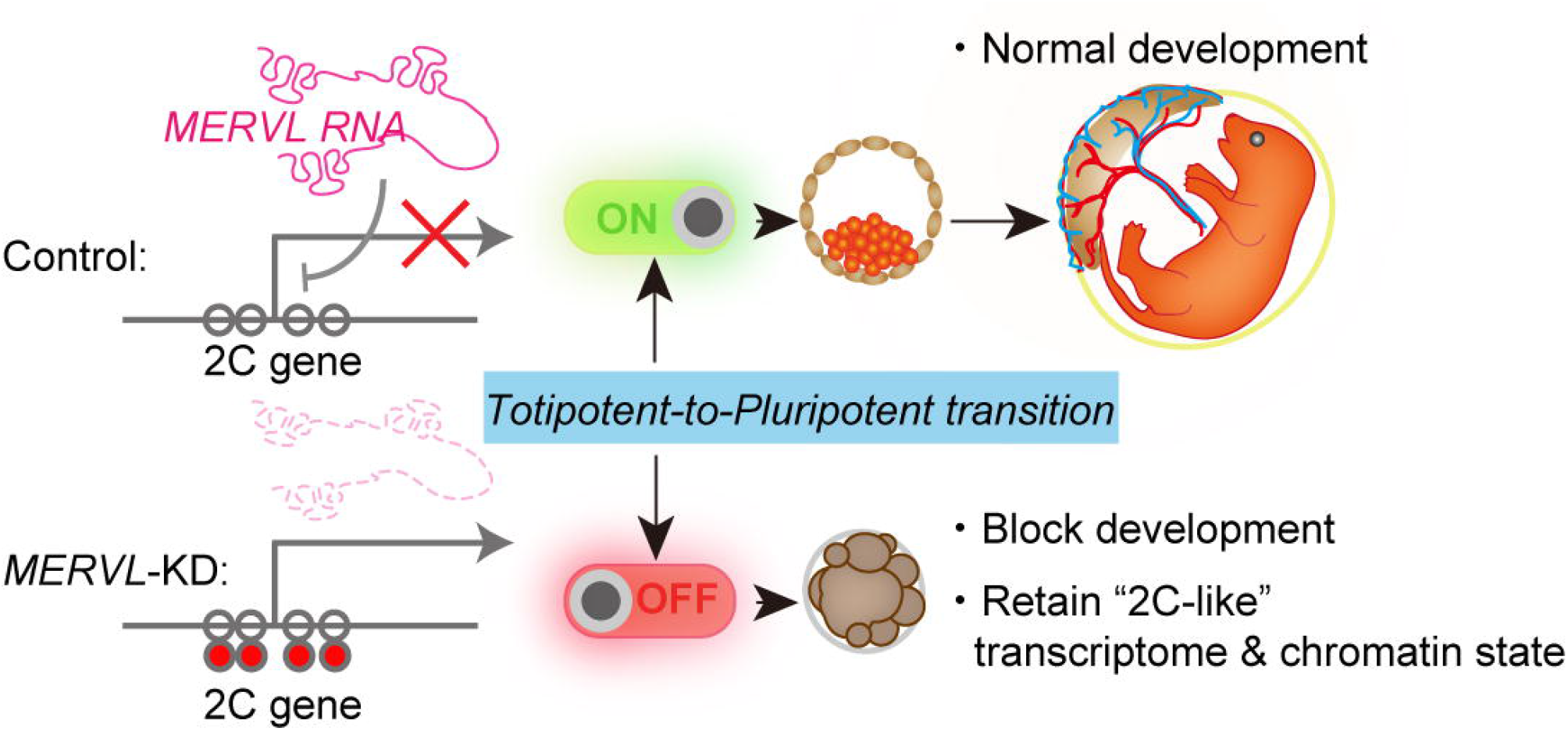
Model: MERVL-mediated totipotent-to-pluripotent transition during preimplantation development. MERVL is expressed in zygote and 2-cell stage-specific manner and may suppresses a subset of 2C genes through an unknown mechanism to facilitate the totipotent-to-pluripotent transition. Knockdown of MERVL RNA compromises these processes and results in embryonic lethality and retention of 2C-like states.

### Differences from previous reports

In this study, one of most striking consequences of loss of MERVL was complete defects in preimplantation development. Until recent past, several groups have used KD strategies in an effort to characterize the role of MERVL in preimplantation development. However, results from these studies were inconsistent. By performing liposome-based transfection with one phosphorothioated ASO against MERVL, Kigami *et al*. have taken up this challenge and reported that MERVL-KD embryos displayed a ∼50% decrease in the developmental competence to the 4-cell stage but has no significant impact on developmental rate to the morula and blastocyst stages, as calculated using the fraction of 4-cell stage embryos that are present ^11^. On the other hand, Huang *et al*. recently performed siRNA-mediated KD against MERVL by microinjection of siRNA into the cytoplasm of zygotes, at two different concentrations (i.e. 20 µM and 80 µM) ^20^. siRNA-mediated KD of MERVL exhibited a mild developmental delay at 4-to 8-cell stage, but no statistically significant differences were observed on developmental rate, morphology, and cell numbers at blastocyst stage between control and KD embryos ^20^. These previous studies found that MERVL may have functional roles in preimplantation development but is not essential for normal development. In sharp contrast, ASO-mediated KD of MERVL in this study results in embryonic lethality with severe defects in first lineage specification and genomic stability, in preimplantation development. The phenotypic differences appear to be explained by methodologies for KD. First, we microinjected ASOs into zygotes resulting in efficient degradation of nuclear MERVL RNA at higher levels than liposome-based transfection (Figure 2b and Figure 3f). Furthermore, it is known that ASOs and siRNA use different mechanisms to degrade RNA targets ^38^. Consistent with previous studies, we confirmed that siRNA-mediated KD of MERVL did not affect development to the blastocyst stage (Figure 3a-d). These results indicate that nuclear MERVL RNA (but not encoded retroviral proteins) is required for progression of preimplantation development. Notably, ASO- and siRNA-mediated MERVL-KD embryos in this study showed no visible signals for either RNA or retroviral proteins as determined by smFISH and immunofluorescence analysis (Figure 2b and Figure 3f), suggesting that our synthesized oligonucleotides efficiently degrade MERVL transcripts (Supplementary Figure S1 and S2). In addition, whereas the activation of *MT2_Mm* inducing the 2C state in previous study is readily appreciable ^16,17^, we also found that the expression levels of chimeric fusion transcripts with *MT2_Mm* were unchanged in MERVL-KD 2-cell stage embryos (Supplementary Figure S4c), suggesting that the phenotypes of MERVL-KD embryos are unlikely to arise from dysregulation of chimeric fusion transcripts with *MT2_Mm*. Overall, to our knowledge, this study is first report to show the importance of MERVL in mouse preimplantation development.

### The regulatory mechanism of MERVL loci in germline

MERVL and its LTR-driven gene transcripts are exclusively transcribed in a 2-cell stage-specific manner. In ESCs, their silencing is mainly mediated by H3K9me2 and the protein complex responsible for its deposition. While other classes of ERVs are bound and repressed by SETDB1 and Kap1/TRIM28 complex, MERVL is highly enriched for G9a-dependent H3K9me2 ^50^. However, the expression level of MERVL was not significantly changed in fully-grown oocytes and 2-cell stage embryos from G9a conditional KO mice ^51^, suggesting that a mechanism other than G9a-dependent H3K9me2 deposition might be responsible for MERVL repression in oocytes and preimplantation. This mechanism remains unknown, a question for future studies.

Although the onset of MERVL expression is accompanied by the minor and major wave of ZGA (Figure 1), key regulators responsible for activation of MERVL loci remain elusive. Recently, it was reported that DUX and ZSCAN4C can directly bind to MERVL loci and robustly drive their expression in the transition from the ESC to 2CLC state, a rare population in ESC cultures ^13,52^. Additionally, *Dppa2* and *Dppa4*, maternally expressed genes present at fertilization, were identified as upstream mediators of murine ZGA, consistent with studies in an ESC model ^53^. However, KO mice of these transcriptional regulators only caused minor defects in preimplantation development and ZGA ^22,23,49,54^. Unlike KO phenotypes for transcriptional regulators, MERVL-KD embryos obviously showed aberrant ectopic transcriptome and chromatin states that resemble the 2C state (Figure 4 and Figure 5) and resulted in embryonic lethality (Figure 2). Furthermore, we recently found that the multicopy homeobox gene *Obox4* can redundantly drive ZGA with Dux (Guo Y., in prep). The KD experiment revealed that single KDs of Obox4 or Dux did not affect preimplantation development, whereas Obox4/Dux double KD completely compromised preimplantation development (Guo Y., in prep). These observations suggest that multiple regulatory networks and axes have mutually compensatory functions, thus securing precise MERVL expression that is required for preimplantation development.

Another recent progress in identifying the upstream regulators of 2C genes has shed light on the DNA damage and p53 activity. Grow *et al*. recently demonstrated that chemically induced DNA damage invoked a DNA damage response (DDR) which in turn led to the activation of p53 followed by 2C genes ^40^. Together with the above report, the present study reveals that MERVL depletion leads to significant upregulation of phosphorylated p53 during preimplantation development (Figure 4j, k). This result is consistent with downregulation of genes related to the “DNA repair pathway” (Supplementary Figure S3c), suggesting that sustained p53 activity in MERVL-KD embryos contributes to retention of a 2C-like transcriptome and chromatin state. This mechanism may also give rise to embryonic lethality due to p53-dependent apoptosis, at the mid-preimplantation stage.

### Predicted function of non-coding MERVL RNA

To investigate the functional essence of MERVL products, we performed siRNA-mediated KD and an RNA rescue assay (Figure 3) and showed that transcription itself and/or cis-acting RNA from MERVL loci is likely to play a critical role for progression of host preimplantation development, rather than through the trans-acting RNA and retroviral protein. Of note, having observed a subset of 2C genes retaining expression in MERVL-KD mid-preimplantation embryos (Figure 4g-i), MERVL RNA may act to repress expression of 2C genes during preimplantation development, at least in part. Although it is unclear how MERVL regulates the transition from totipotency to pluripotency, several lines of evidence allow us to predict the mechanism by which MERVL regulates host transcriptome and chromatin state in preimplantation development. First, a recent study demonstrated that MERVL drives the 3D reorganization of the genome in the 2CLC and early mouse embryos, by providing a topologically associating domain (TAD) boundary that is coupled to directional transcription from MERVL ^55^(Kruse *et al*., 2019 preprint). Thus, it is plausible that transcription itself from MERVL loci might be responsible for remodeling of chromatin structure during the transition from totipotency to pluripotency. Alternatively, MERVL *cis*-acting RNA may have important roles in regulation of the host genome (that are not captured by the *trans*-acting RNA rescue assay (Figure. 3e-h)). MERVL is an intronless gene, a key feature of *cis-*acting RNAs, because poor-or un-spliced non-coding RNA is tethered to chromatin in an U1 small nuclear ribonucleoprotein-dependent manner and promotes the formation of spatial compartments in the nucleus ^56^. Curiously, the expression of PRC2-target genes were upregulated in MERVL-KD embryos (Figure 4f), implying that deposition of the repressive H3K27me3 mark is attenuated through the genome. Previous studies demonstrated that *Xist*, a representative non-coding RNA, acted in *cis-*, to recruit PRC2 and initiate X chromosome inactivation ^57^. Likewise, there is another possibility that MERVL transcript may act as a non-coding RNA, to recruit PRC2 and/or other epigenetic regulator to target loci. Future studies can now interrogate the intriguing mechanisms that may underlie the important role of MERVL transcript in regulating the host genome we have uncovered here.

## Methods

### Animals

Eight-week old female and male B6D2F1 (BDF1) mice were purchased from the Japan SLC, Inc. All animal experiments were reviewed and approved by the Institutional Animal Care and Use Committee (protocol no. 09105-(10) and 11045-(6)) at Keio University, where mice were maintained and fed *ad libitum* with standard diet and water in a temperature- and light-controlled room (23°C±3°C, 14 light/10 dark cycle).

### Cell culture

Doxycycline (Dox)-inducible Dux ESC line was generated using the PiggyBac Transposon System in our laboratory (hereafter referred to as ESC^DUX^) (Guo Y., in prep), in which Dux expression was induced in the presence of Dox. The ESC^DUX^ was cultured in ESC medium (15% FBS, 25 mM HEPES, 1× GlutaMAX, 1× MEM non-essential amino acids solution, 1× penicillin/streptomycin and 0.055 mM β-mercaptoethanol in DMEM high glucose (4.5 g l^-1^)) containing 2i (1 µM PD0325901, LC Laboratories; and 3 µM CHIR99021, LC Laboratories) and LIF (1,300 U ml^-1^, in-house) on cell culture plates coated with 0.2% gelatin under feeder-free conditions. The expanded colonies were dissociated using 0.25% trypsin-EDTA solution for passaging.

### MERVL-targeting ASO design

Three independent ASOs were designed based on the consensus sequence of MERVL (Supplementary DataSet5). ASO candidates that effectively target the internal retroviral regions of MERVL were selected using ChIRP Probe Designer (https://www.biosearchtech.com/support/tools/design-software/chirp-probe-designer), taking into account the secondary structure of the MERVL RNA. To identify potential binding sites for MERVL-targeting ASOs *in silico*, non-redundant highly confident MERVL loci were downloaded from RepeatMasker database (http://www.repeatmasker.org/species/mm.html) and converted to BED format. Consequently, FASTA files for these MERVL loci were extracted using the getfasta function as implemented in BEDTools (version 2.30.0)^58^ and input to makeblastdb program from Blastn (version 2.6.0+) to generate DNA database for BLAST search. Each ASO sequence was used to search the database of MERVL DNA using blastn ^59^, with up to two mismatches allowed. The three independent ASOs against MERVL with the highest targeting rate were used for knockdown experiments (Supplementary DataSet5).

### ASO-mediated KD of MERVL in ESCs

A total of 1 × 10^5^ ESC^Dux^s were harvested from culture dishes per one nucleofection. To validate efficiency of the ASO-mediated KD, the cells were nucleofected with 1 nmol ASO against MERVL or Scramble ASO using P3 Primary Cell 96-well Nucleofector Kit (Lonza) according to manufacturer’s instruction and seeded into each well of a 24-well plate containing 500 µl of ESC medium with 2i, Lif and 1.1 µl of iMatrix-511 Silk solution (0.5 mg/ml, Nippi). After 6 hrs. of nucleofection, we added 0.5 µl of 10 µg ml^-1^ Dox to each well of 24-well plate. The following day (24 hrs. post-nucleofection), the cells were harvested for quantitative RT-PCR and Western blotting to evaluate the expression of MERVL.

### Embryo culture and microinjection

Female mice were superovulated by intraperitoneal administrations of 150 µl of CARD HyperOva (Kudo) followed 48 hrs. later with 7.5 IU of human chorionic gonadotropin (hCG, Asuka Pharmaceutical Co.). After injection of hCG, females were housed with male mice overnight for copulation. The following day, zygotes were collected from the oviducts of superovulated/mated females. The microinjection was carried out under a phase-contrast inverted microscope (IX73, Olympus) equipped with a micromanipulation system (Narishige). Each ASO (20 µM) and siRNA (20 and 80 µM) was microinjected into the male pronuclei of zygotes using FemtoJet 4i (Eppendorf). All ASO and siRNA sequences used in this study are listed in Supplementary Dataset 5. Embryos were cultured in KSOM (MR-101-D, Sigma) at 37°C with 5% CO_2_.

### RNA rescue assay

ASO-resistant full-length MERVL was constructed by substituting all of the three ASO-targeted sequences of MERVL using inverse PCR. The primers used to produce the constructs are listed in Supplementary Dataset 5. Briefly, intact MERVL sequence was obtained from a bacterial artificial chromosome (BAC) plasmid (RP23-231A8, https://www.ncbi.nlm.nih.gov/nuccore/AC127321.4, Thermo Fisher Scientific) and subcloned into pCAGGS_Myc plasmid using NEBuilder HiFi DNA Assembly Master Mix (New England Biolabs). Individual ASO-targeted sequences were mutated by inverse PCR with PrimeSTAR GXL DNA polymerase (TaKaRa) and specific primer sets (Supplementary Dataset 5). For subsequent steps, ASO-resistant full length MERVL was amplified by PCR with a specific primer set containing the T7 promoter sequence at the 5’ end of forward primer for *in vitro* transcription (IVT) with T7 polymerase. Following PCR amplification, IVT was performed using MEGAscript T7 Transcription Kit (Thermo Fisher Scientific) according to the manufactures’ instruction. Transcribed RNA was purified using RNeasy Mini Kit (QIAGEN). ASO-resistant MERVL RNA was co-microinjected into the male pronucleus of zygotes with 20 µM ASOs (mixed) at the final concentration of 200 ng µl^-1^.

### smFISH and immunofluorescence analysis

Embryos were fixed using fixative solution (4% paraformaldehyde and 0.2% polyvinyl alcohol (PVA) in PBS) for 20 min at RT, then permeabilized with 0.5% TritonX-100 in PBS for 10 min at RT. For smFISH, embryos were washed once with Stellaris Wash Buffer A (Bioresearch technologies) for 5 min at RT. FISH was then performed overnight at 37°C in Stellaris Hybridization Buffer (Bioresearch technologies) containing 25 pmol of 48 probes set (Quasar 670-labeled, Bioresearch technologies) against MERVL RNA. The following day, embryos were washed once with Stellaris Wash Buffer A for 30 min at 37°C and counterstained with DAPI (1 µg ml^-1^, Nacalai Tesque) for 30 min at 37°C. After washing with Stellaris Wash Buffer B (Bioresearch technologies) for 5 min at RT, embryos were transferred to 10 µl drops of 0.2% PVA in PBS on a glass-bottomed dish covered by paraffin oil. The FISH probe sequences used in this study are listed in Supplementary Dataset 5. For immunofluorescence analysis, non-specific immunoreaction was blocked by incubating embryos in 2% bovine serum albumin (BSA) in PBS for 1 hr. at RT. After blocking, whole-mount immunofluorescence analyses were conducted using the following primary antibodies: mouse anti-MERVL Gag (1/5 dilution, kept in our lab), rabbit anti-OCT4 (1/100 dilution, ab181557, abcam), mouse anti-CDX2 (1/100 dilution, ab157524, abcam), mouse anti-E-Cadherin (1/100 dilution, 610182, BD Transduction Laboratories), rabbit anti-Cleaved Caspase-3 (1/200, 9661, CST) and rabbit anti-Phospho-S15-p53 (1/200 dilution, 9284, CST). Embryos were incubated overnight at 4°C with primary antibody and then washed three times with 2% BSA in PBS for 5 min each and incubated with Alexa Fluor 488- and/or 568-conjugated anti-mouse and/or anti-rabbit IgG secondary antibodies (1/500, Thermo Fisher Scientific) for 1 hr at RT. After washing three times with 0.2% BSA in PBS for 10 min each, the embryos were counterstained with DAPI (1 µg ml^-1^) for 30 min at RT and transferred to 10 µl drops of 0.2% PVA in PBS on a glass-bottomed dish covered by paraffin oil. smFISH and immunofluorescence images were obtained with a BZ-X810 fluorescence microscope (Keyence) or FV3000 confocal laser scanning microscope (Olympus) and processed with ImageJ (NIH).

### EU (5-Ethynyl Uridine) incorporation assay

The embryos microinjected with either scramble ASOs or MERVL ASOs, were cultured in KSOM for 24 hrs. On the next day, 2-cell stage embryos were transferred to new KSOM containing 1 mM 5-ethynyl uridine (EU, from Click-iT RNA Alexa Fluor 488 Imaging Kit, Thermo Fisher Scientific) and cultured for 1h at 37°C with 5% CO_2_. The EU-labeled embryos were immediately fixed and permeabilized as described above. Incorporated EU into newly synthesized RNA was detected using the Click-iT RNA Alexa Fluor 488 Imaging Kit (Thermo Fisher Scientific) according to the manufacturer’s instructions. In brief, the fixed embryos were treated with the Click-iT reaction cocktail containing Alexa Fluor 488 azide for 30 min at RT, and then washed once with Click-iT reaction rinse buffer. After washing once with 0.2% PVA in PBS, the embryos were counterstained with Hoechst 33342 (2 µg ml^-1^, Dojindo) for 30 min at RT and transferred to 10 µl drops of 0.2% PVA in PBS on a glass-bottomed dish covered by paraffin oil. Images of EU-labeled embryos were obtained with a FV3000 confocal laser scanning microscope (Olympus) and processed with ImageJ (NIH).

### Quantitative RT-PCR

Acidic Tyrode’s solution (Sigma) containing 0.2% PVA was used to remove the zona pellucida (ZP) from embryos before sample collection. 30 ZP-free embryos and nucleofected ESC^Dux^s were lysed in Lysis Solution (from SuperPrep II Cell Lysis & RT Kit, TOYOBO), directly. Genomic DNA elimination and reverse transcription were performed using the SuperPrep II Cell Lysis & RT Kit with random hexamers and oligo dT primers according to the manufacturer’s instruction. Real-time quantitative PCR was carried out using the following conditions: 95°C for 1 min, followed by 45 cycles each of 95°C for 15 sec and 60°C for 1 min, on a Thermal Cycler Dice Real Time System III (TaKaRa) with Thunderbird SYBR qPCR Mix (TOYOBO) and specific primer sets (Supplementary Dataset 5). Relative gene expression was quantified with the ΔΔCT method and normalized to *Actb* or *Gapdh* expression.

### Western blotting

The cells were directly lysed in Laemmli SDS sample buffer (1x, 62.5 mM Tris-HCl pH6.8, 2% SDS, 10% Glycerol, 5% β-Mercaptoethanol and 0.02% Bromophenol blue) and sonicate with Bioruptor at High setting, for 10 cycles each of 30 s with 30 s intervals. Total proteins were denatured at 95°C for 5 min and separated by 10% SDS-PAGE. The proteins were then transferred onto a Protran Nitrocellulose Membranes (0.45 µm pore size, GE Healthcare) via Power Blotter-Semi-dry Transfer System (Thermo Fisher Scientific). The membrane was blocked with Bullet Blocking One for Western Blotting (Nacalai Tesque) for 10 min at RT with gently rocking before incubation with mouse anti-MERVL Gag (1/5 dilution, kept in our lab) or mouse anti-β-tubulin (1/2000 dilution, kept in our lab) antibody in 1:20 diluted blocking buffer with 0.1% PBST at 4°C overnight. The following day the membranes were then incubated with HRP-conjugated goat anti-mouse IgG secondary antibody (1/10000 dilution, #330, MBL life science) in 1:20 diluted blocking buffer with 0.1% PBST for 30 min at RT with gently rocking. After washing three times with 0.1% PBST for 10 min each, Blots were developed using ECL Western Blotting Detection Reagent (Sigma) and exposed onto X-ray film.

### RNA-seq

Preparation of total RNA-seq libraries was performed using SMART-Seq Stranded Kit (Clontech), according to the manufacturer’s instructions. In brief, 30 ZP-free embryos were lysed in 1x Lysis Buffer containing RNase inhibitor (0.2 IU/µl, from SMART-Seq Stranded Kit, Clontech), directly. RNAs were randomly sheared by heating at 85°C for 8 min and subjected to reverse transcription with random hexamers and PCR amplification. Ribosomal fragments were depleted from each cDNA sample with scZapR and scR-Probes. Indexed total RNA-seq libraries were enriched through a second PCR amplification and sequenced using an Illumina HiSeqX sequencer (paired-end, 150 bp). Two biological replicates were generated for each sample.

### miniATAC-seq

The protocol for miniATAC-seq library was adapted from a previous report with minor modifications ^45^. Briefly, 30 ZP-free embryos were lysed in 6 µl of Lysis Buffer (10 mM Tris-HCl (pH7.4), 10 mM NaCl, 3 mM MgCl_2_ and 0.5% NP-40) for 10 min on ice. After cell lysis, 2 µl of ddH_2_O, 10 µl of 2x TD Buffer (from Nextera XT DNA Prep Kit, Illumina) and 2 µl of Tn5 Transposase (from Nextera XT DNA Prep Kit, Illumina) were added to the lysates, which were mixed by pipetting followed by incubation at 37°C For 30 min. To stop the transposase reaction, 5 µl of 0.5% SDS was added and incubated at RT for 5 min. After Tn5 tagmentation, 100 ng Carrier RNA (Yeast tRNA, Sigma) was added and then messed samples up to 130 µl with 1x TE Buffer (10 mM Tris-HCl (pH8.0) and 1mM EDTA). Tagmented DNA was purified by phenol-chloroform extraction and ethanol precipitation with 20 µg of glycogen as carrier and dissolved in 20 µl of ddH_2_O. Afterwards, PCR was performed to amplify and index libraries using the following conditions: 72°C for 5 min and 98°C for 30 sec, followed by 18 cycles each of 98°C for 10 sec, 63°C for 30 sec and 72°C for 1 min, with NEBNext HiFi PCR Master Mix and specific index primer sets (Supplementary Dataset 5). Pooled miniATAC-seq libraries were sequenced using an Illumina HiSeqX sequencer (paired-end, 150 bp). Two biological replicates were generated for each sample.

### RNA-seq and ATAC-seq data processing

Methods for RNA-seq analysis have been described previously ^60^, In short, raw paired-end RNA-seq reads were aligned to indexed mouse genome (GRCm38/mm10) using STAR aligner version 2.5.3a ^61^ with following options, --twopassMode Basic; -- outSAMtype BAM SortedByCoordinate; --outFilterType BySJout; -- outFilterMultimapNmax 1; --winAnchorMultimapNmax 50; --chimSegmentMin 12; -- chimJunctionOverhangMin 8; --alignSJoverhangMin 8; --alignSJDBoverhangMin 10; -- outFilterMismatchNmax 999; --outFilterMismatchNoverReadLmax 0.04; -- alignSJstitchMismatchNmax 5 -1 5 5; --outSAMattrRGline ID:GRPundef; -- alignIntronMin 20; --alignIntronMax 1000000; --alignMatesGapMax 1000000 for unique alignments. To quantify aligned reads on annotated genes (Genecode vM25) and repetitive loci (originating from mm10.fa.out (RepeatMasker), best match TE annotation (defined in ^60^, we used featureCounts function, part of the Subread package ^62^. To detect differentially expressed genes and repetitive elements between control and MERVL-KD embryos, a output file of featureCounts was input to DESeq2 package (version 1.16.1); then, the program functions DESeqDataSetFromMatrix and DESeq were used to compare each gene’s expression level between two biological samples. Differentially expressed genes were identified through two criteria: (1) ≥2-fold change and (2) binominal tests (*Padj* < 0.01; P values were adjusted for multiple testing using the Benjamini-Hochberg method). To perform GO and ChEA analyses, we used the functional annotation clustering tool in Enrichr ^63^. The annotated terms with *P* < 0.05 from a modified Fisher’s exact test were considered significant. To visualize read enrichments over representative genomic loci, TDF files were created from sorted BAM files using the IGVTools count function (Broad Institute). Figures of continuous tag counts over selected genomic intervals were created in the IGV browser (Broad Institute).

Raw paired-end ATAC-seq reads were filtered by TrimGalore (version 0.6.4) with the default setting and aligned to indexed mouse genome (GRCm38/mm10) using bowtie2 (version 2.4.4) ^64^ with the following options, -N 1; -L 25; --no-mixed; --no-discordant. Multiple aligned reads and PCR duplicates were removed using grep -v ‘XS:’ and MarkDuplicates function with REMOVE_DUPLICATES=true option, a part of Picard tools. Using SeqMonk (Barbraham Bioinformatics), we calculated Pearson correlation coefficients between biological replicates. Peak calling for ATAC-seq data was performed using MACS2 (version 2.1.4)^65^ with default arguments; we used a cut-off of P ≤ 10^−5^. The average tag density plots were drawn using Ngsplot (version 2.47.1) and plotHeatmap program as implemented in deepTools (version 3.1.3)^66^. To visualize read enrichment over representative genomic loci, TDF files were created from sorted BAM files using the IGVTools count function (Broad Institute). Figures for continuous tag counts over selected genomic intervals were created in the IGV browser (Broad Institute). For k-means clustering of ATAC-seq peaks, we firstly generated a set of regions that were called peaks with MACS2 in at least one of the samples, by merging ATAC-seq peak regions of all biological replicates using the mergeBed function from BEDTools (version 2.30.0). Using this merged peak file and ATAC-seq data from control embryos, we ran k-means clustering analysis using computeMatrix and plotHeatmap program as implemented in deepTools (version 3.1.3)^66^ and determined that k=5 was suitable for our data. To evaluate the functional annotation of each cluster, we utilized Genomic Region Enrichment of Annotation Tool (GREAT, version 4.0.4)^67^ and HOMER (version 4.9)^68^, which associates ATAC-seq peaks in each cluster with their genomic feature and ontology of their putative target genes adjacent to peaks.

## Data availability

The total RNA-seq and miniATAC-seq data from control and MERVL-KD embryos are deposited in the Gene Expression Omnibus (GEO) under accession code GSE196520 (Token number to access private data is gpufgwswldepzej).

## Author contributions

The manuscript was written by A.S and H.S., with critical feedback from all other authors, A.S., H.I. and H.S conceived and designed this study. K.M. provided instruction for biochemistry and molecular biology. H.I. developed ASOs against MERVL. T.K. and A.S performed microinjection for generating MERVL-KD embryos and RNA rescue assay with the help of H.M. and carried out phenotypic analyses of generated MERVL-KD embryos. A.S. and T.K. performed total RNA-seq and ATAC-seq analyses. T.K., Y.G. and A.S designed and interpreted bioinformatics analyses with the help of M.A. A.S. and H.S supervised the project.

## Acknowledgements

We thank all members of the Siomi lab for discussions on this work, Dr. Katsuhiko Hayashi (Graduate School of Medicine, Faculty of Medicine, Osaka University) and Dr. Azusa Inoue (Center for Integrative Medical Science, RIKEN) for critical reading of the manuscript. This work was supported by JSPS Grants-in-Aid for Early-Career Scientists (21K15108 to A.S.), the Kato Memorial Bioscience Foundation Research Grant (to A.S.), MEXT Grants-in-Aid for Scientific Research in Innovative Areas (19H05753 to H.S.), AMED project for elucidating and controlling mechanisms of aging and longevity (1005442 to H.S.), the Uehara Memorial Foundation Research Incentive Grant (to A.S.), and the Uehara Memorial Foundation Research Grant (to H.S.).

## Supplementary Figure legends

**Figure S1. Design and validation of ASOs targeting MERVL**.

**(a)** Schematic of MERVL indicating the positions of ASOs. Below whisker plot showing alignment quality in each interspersed genomic MERVL copy (adapted from Dfam: https://dfam.org/family/DF0003918/seed). **(b)** Pie chart indicates the populations of full-length (≥ 5001 bp) and truncated (1-5000 bp) MERVL copies across the mouse genome. **(c)** Chromosome maps showing the distribution of MERVL copies that are targeted by ASOs throughout the mouse genome. Each colored rectangle (blue for ASO-#1, yellow for ASO-#2, and red for ASO-#3) indicates targeted MERVL copy. **(d)** Histogram showing the distributions of ASO-targeted (magenta) and untargeted (green) MERVL copies. The abundance of each length of MERVL element was tallied with a given count. The proportion of full length MERVL (>5 kb) was highlighted in red. **(e)** Schematic for ASO-mediated KD against MERVL in ESC harboring a doxycycline (Dox)-inducible *Dux* transgene (termed ESC^DUX^). TRE, tetracycline responsive element; rtTA, reverse tetracycline responsive transcriptional activator. **(f)** Expression levels of MERVL mRNA measured by quantitative RT-PCR in ESC^DUX^s nucleofected with each individual MERVL-targeting ASO and Scramble ASO. Relative expression is quantified with ΔΔCt method and normalized to *Gapdh* expression. Error bars represent mean± s.e.m. **(g)** Expression level of MERVL mRNA measured by quantitative RT-PCR in control and ASOs (Mixed)-mediated MERVL-KD ESC^DUX^s in the presence or absence of Dox. Relative expression is quantified with ΔΔCt method and normalized to *Gapdh* expression. Error bars represent mean± s.e.m. **(h)** Expressions of MERVL retroviral protein (Gag), measured by western blotting in control and ASOs (Mixed)-mediated MERVL-KD ESC^DUX^s in the presence or absence of Dox. β-Tubulin was used as a loading control.

**Figure S2. The KD effectiveness of each ASO against MERVL in preimplantation embryos**.

**(a)** Representative images of smFISH with probes set for full-length MERVL RNA sequence and DNA counterstained with DAPI in 2-cell stage embryos from each experimental condition. Scale bar: 20 µm. **(b)** Representative phase-contrast Images of 4.5 dpc blastocysts from each experimental condition. Scale bar: 100 µm. **(c)** Percentage of embryos by developmental stage at 4.5 dpc in each experimental condition.

**Figure S3. Total RNA analysis of control and MERVL-KD embryos at 2-cell, 4-cell, and 8-cell stages**.

**(a)** Schematic of sample collection for total RNA-seq analysis in control and MERVL-KD embryos. We supplied the apparently healthiest embryos upon MERVL-KD for preparation of a total RNA-seq library. **(b)** Scatter plots showing the reproducibility between biological replicates in total RNA-seq data. Pearson correlation values (R) are shown. **(c)** GO analysis of DEGs in MERVL-KD embryos, assessed by the Enrichr. The known GO terms associated with cell cycle, DNA repair and apoptosis/cell death are highlighted in green, blue and red, respectively.

**Figure S4. MERVL-KD only dysregulates a few types of transposable element**.

**(a)** Dot plots showing RPKM values for *MERVL-int* and *MT2_Mm* in control and MERVL-KD embryos at 2-cell, 4-cell, and 8-cell stages. **(b)** MA plots show differentially expressed (DE) repetitive elements (annotated in RepeatMasker) between control and MERVL-KD embryos at 2-cell, 4-cell, and 8-cell stages. DE repeats were defined as those with a FoldChange≥|2| and *Padj*<0.05 (binomial test with Benjamini–Hochberg correction) using DESeq2 and shown in red circles. *MERVL-int* and *MT2_Mm* were included in downregulated genes. **(c)** Violin plot indicates RPKM values for chimeric fusion transcripts in control and MERVL-KD 2-cell stage embryos. N.S., not significant; two-tailed unpaired t-tests.

**Figure S5. miniATAC-seq analysis of control and MERVL-KD embryos at 2-cell, 4-cell, and 8-cell stages**.

**(a)** Schematic of sample collection for miniATAC-seq analysis in control and MERVL-KD embryos. We supplied apparently healthiest embryos upon MERVL-KD for preparation of miniATAC-seq library. **(b)** Scatter plots showing the reproducibility between biological replicates in miniATAC-seq data. Pearson correlation values (R) are shown. **(c)** Track view of ATAC-seq read enrichments at a representative chromosomal position in all biological replicates.

**Supplementary Dataset 1**

Alignment and quantification statistics in each NGS sample.

**Supplementary Dataset 2**

Transcript profiles of genes and repetitive elements for control and MERVL-KD embryos from total RNA-seq analysis.

**Supplementary Dataset 3**

List of DEGs between control and MERVL-KD embryos.

**Supplementary Dataset 4**

List of locations of ATAC-seq peaks in each cluster and their biological features.

**Supplementary Dataset 5**

List of sequence information used in this study.

