## Supplementary figures and images for "Transcription of Murine Endogenous Retrovirus MERVL Is Required for Progression of Development in Early Preimplantation Embryos"

### Supplementary Figure S1

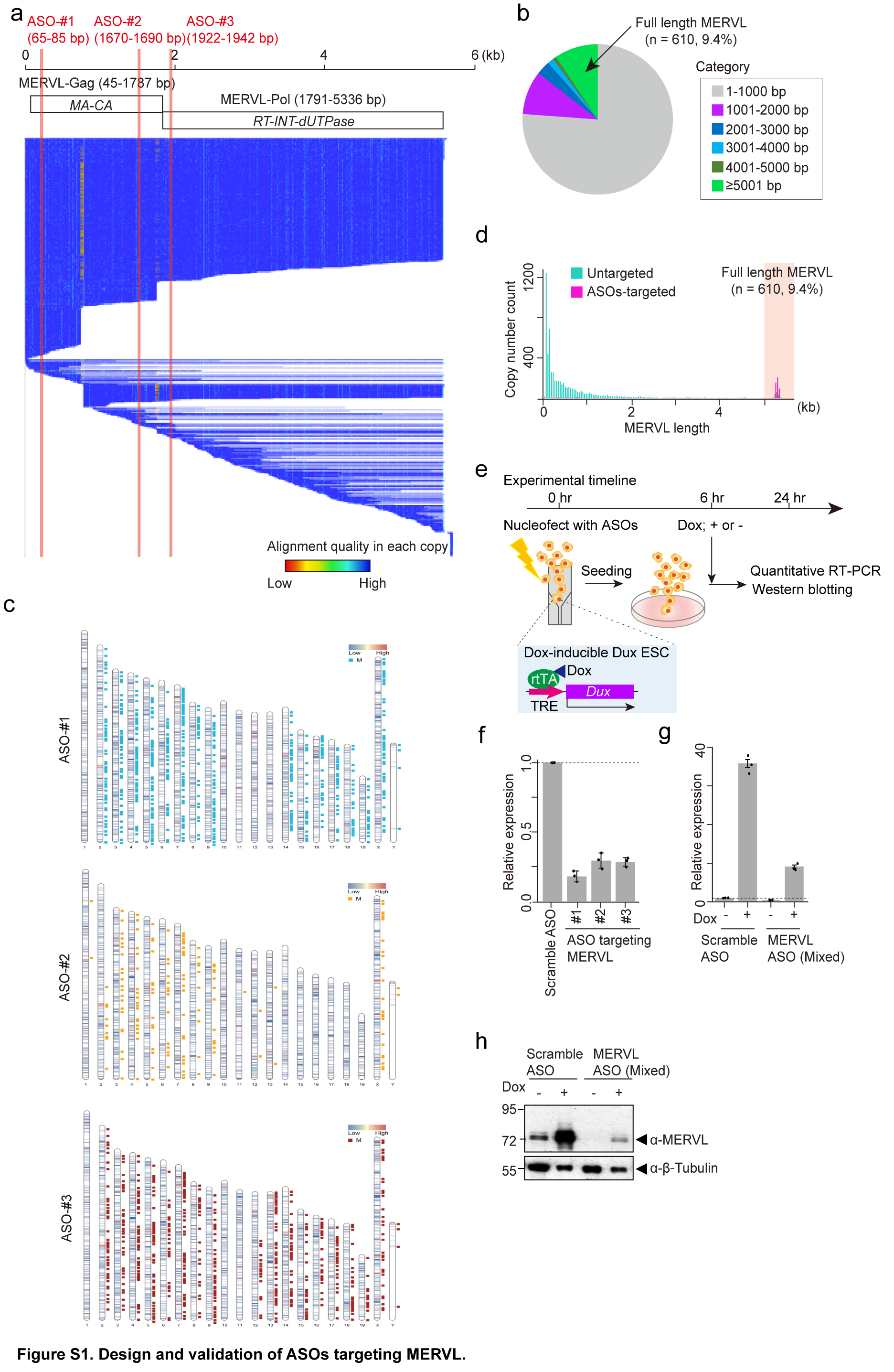

### Supplementary Figure S2

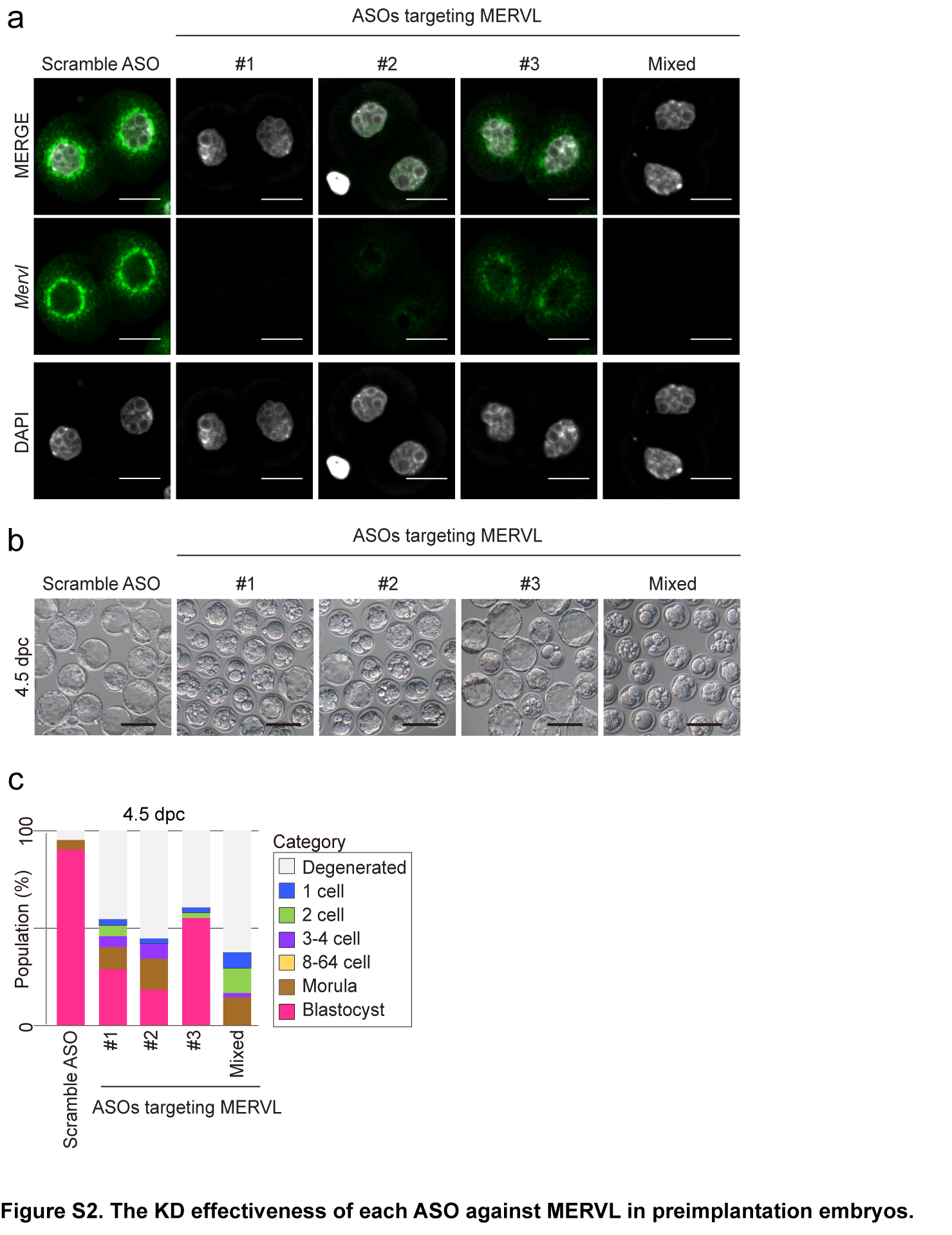

### Supplementary Figure S3

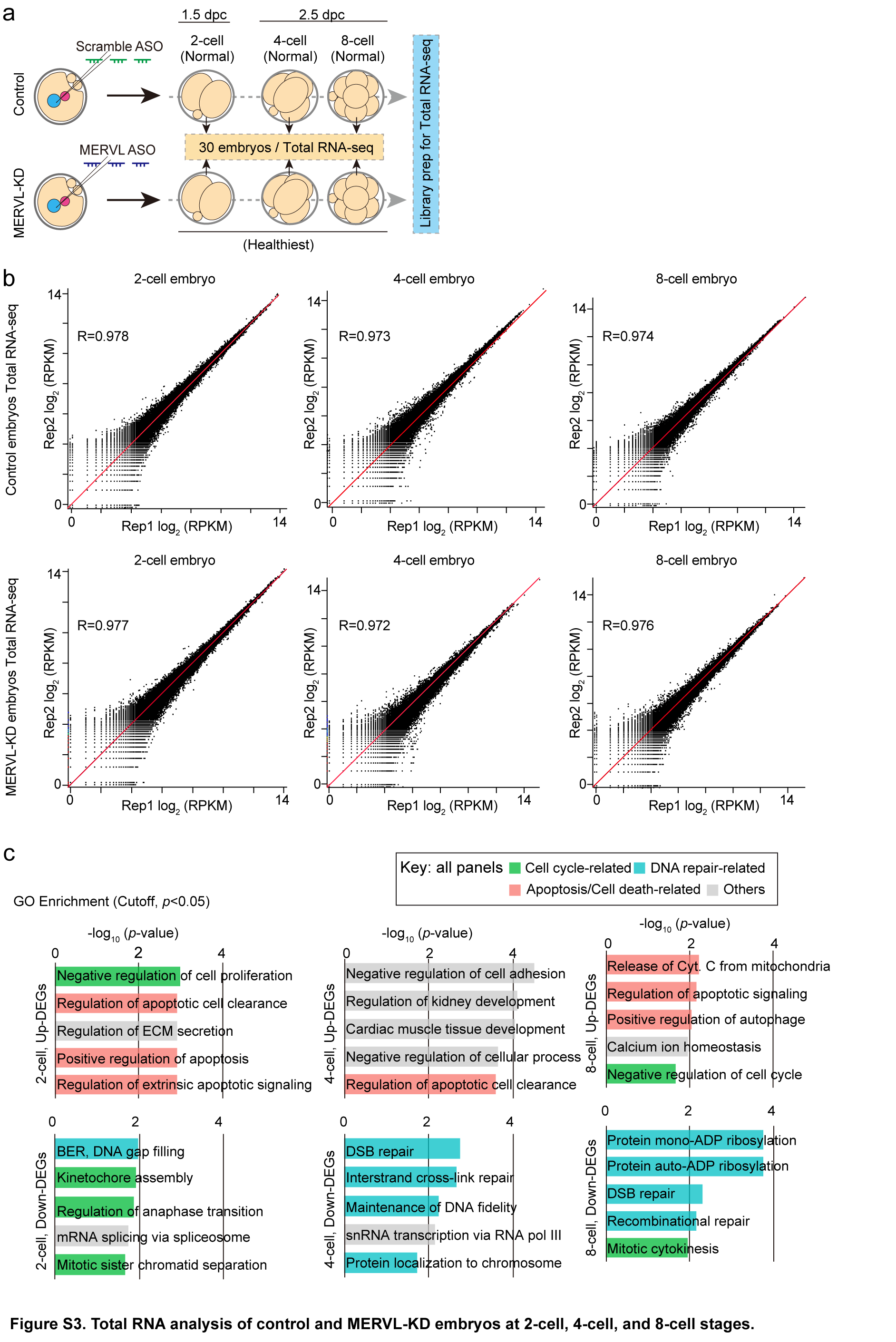

### Supplementary Figure S4

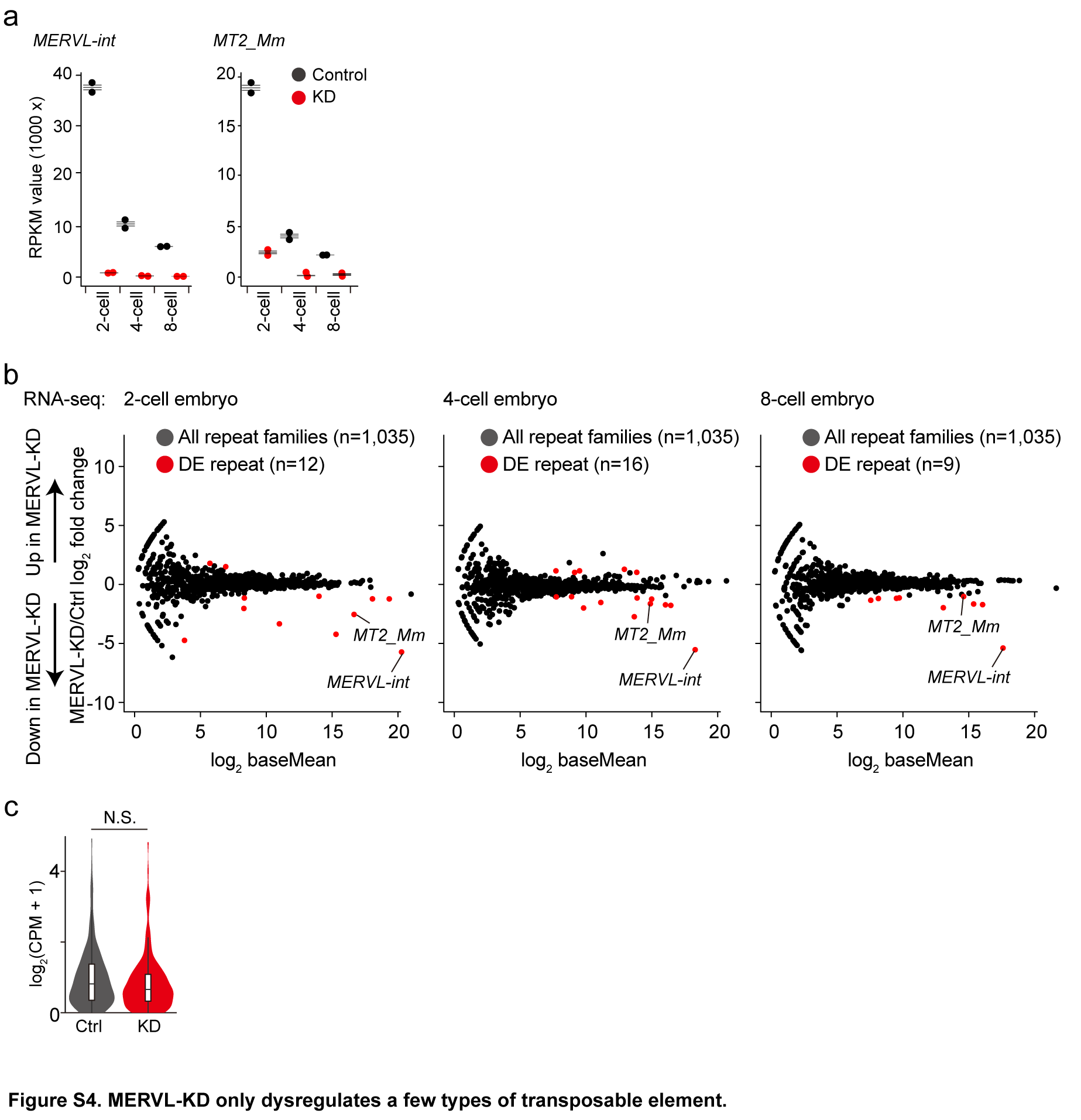

### Supplementary Figure S5

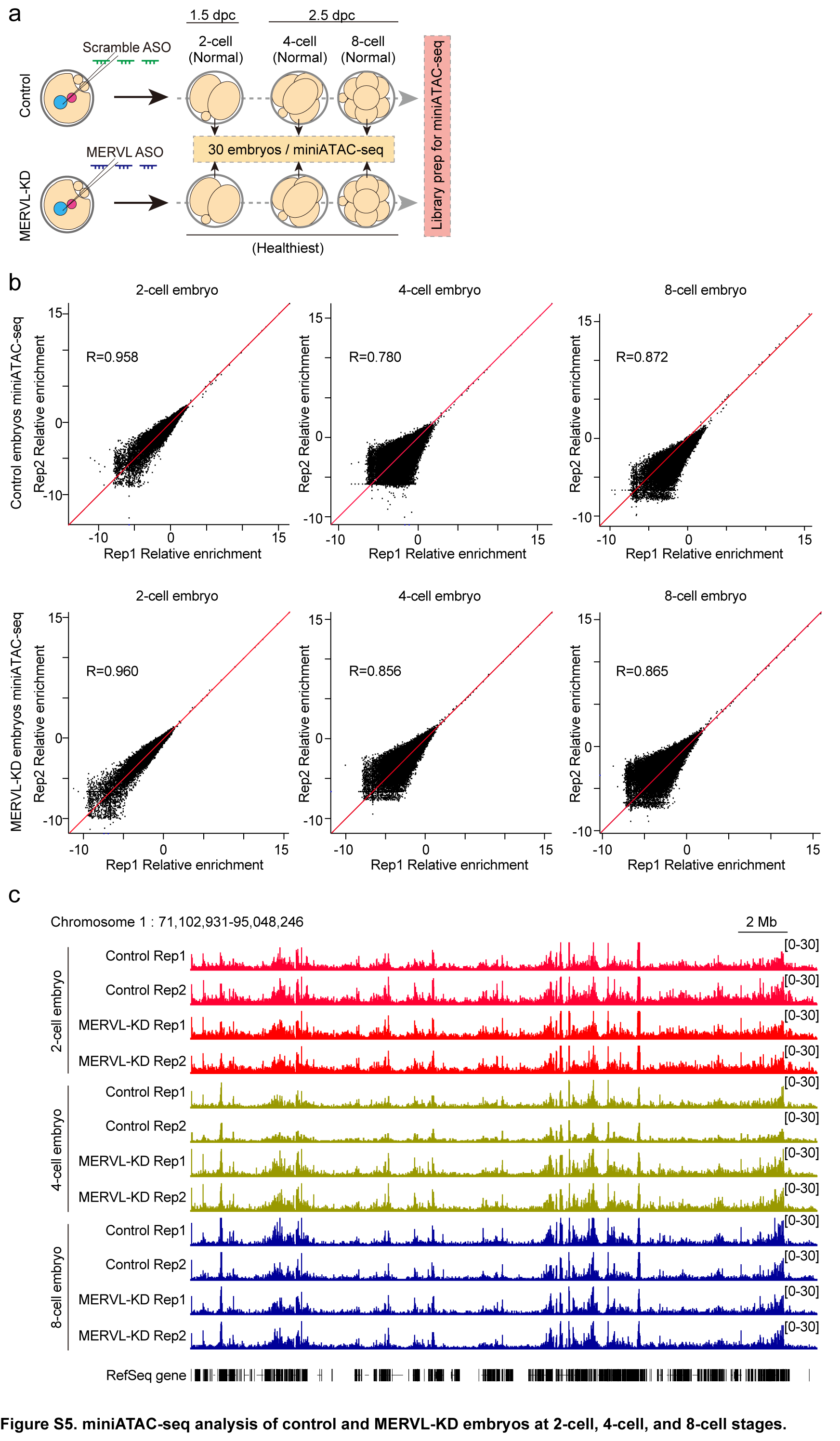
